# Learning temporal integration from internal feedback

**DOI:** 10.1101/2019.12.29.890509

**Authors:** Erik Nygren, Alexandro Ramirez, Brandon McMahan, Emre Aksay, Walter Senn

## Abstract

There has been much focus on the mechanisms of temporal integration, but little on how circuits learn to integrate. In the adult oculomotor system, where a neural integrator maintains fixations, changes in integration dynamics can be driven by visual error signals. However, we show through dark-rearing experiments that visual inputs are not necessary for initial integrator development. We therefore propose a vision-independent learning mechanism whereby a recurrent network learns to integrate via a ‘teaching’ signal formed by low-pass filtered feedback of its population activity. The key is the segregation of local recurrent inputs onto a dendritic compartment and teaching inputs onto a somatic compartment of an integrator neuron. Model instantiation for oculomotor control shows how a self-corrective teaching signal through the cerebellum can generate an integrator with both the dynamical and tuning properties necessary to drive eye muscles and maintain gaze angle. This bootstrap learning paradigm may be relevant for development and plasticity of temporal integration more generally.

**Highlights:**

- A neuronal architecture that learns to integrate saccadic commands for eye position.
- Learning is based on the recurrent dendritic prediction of somatic teaching signals.
- Experiment and model show that no visual feedback is required for initial integrator learning.
- Cerebellum is an internal teacher for motor nuclei and integrator population.

## 1 Introduction

Neuronal integrators are involved in various cortical and subcortical processes. For example, when our eyes scan over a text, when we memorize what we read, and when we make a decision upon the reading, neurons need to accumulate information in time. During scanning eye movements, command signals encoding the velocity of fast eye movements termed saccades are integrated by neurons to generate position signals which fixate the gaze upon a point of interest (Aksay, Olasagasti, et al. 2007). In working memory, a new item is added when the activity of memory neurons incrementally grows upon presentation of a brief stimulus (Goldman-Rakic 1995; Pereira and Brunel 2018). In decision making, continuously presented evidence is gradually accumulated by a neuronal population displaying ramp-like changes in activity prior to the decision (Gold and Shadlen 2007; Brody and Hanks 2016).

A hallmark of neuronal integration is that the underlying network has an integration time constant that is orders of magnitude longer than the membrane time constant of the individual neurons. A range of mechanisms for generating long integration time constants have been proposed, including synaptic facilitation, active dendritic properties, and recurrent network interactions (Seung 1996; Koulakov, Raghavachari, et al. 2002; Goldman et al. 2003; Lim and Goldman 2013; Sanders et al. 2013). All models based on these mechanisms require some degree of tuning to show stable integration, and the tuning requirements tend to be more severe the greater the dependence on recurrent interactions. Since these recurrent interaction models generally have the most explanatory power across the various settings where neural integration has been observed, it is especially important to consider how such networks might learn to perform their tasks.

Here we investigate in the context of oculomotor control the problem of how a neural integrator might develop and autonomously be tuned. Temporal integration in the oculomotor system is important for maintaining the desired gaze angle. The eye position command signals necessary for maintaining gaze are generated in the velocity-to-position neural integrator (VPNI), a hindbrain neuronal population that receives velocity-encoding command signals from vestibular neurons for smooth eye movements such as the vestibulo-oculo reflex, and from burst generator neurons for saccades and fixations. Studies of the mechanism of integration in this system implicate a prominent role for recurrent (excitatory) network interactions in establishing positive feedback: structure-function studies demonstrate local collaterals within the network (Lee, Arrenberg, and Aksay 2015; Aksay, Gamkrelidze, et al. 2001; Vishwanathan et al. 2017), intracellular recordings during gaze holding show that VPNI neurons receive synaptic input consistent with positive feedback (Aksay, Gamkrelidze, et al. 2001), and localized disruption of this feedback leads to expected deficits in the capacity of the network to integrate (Aksay, Olasagasti, et al. 2007). This network is also tunable, with learning of new integration dynamics occurring on the tens of minutes time scale when visual error signals are presented (Major et al. 2004). It is unclear, however, if visual error signals are the only trigger for integrator tuning, or if visual signals are even relevant for the early development of the VPNI.

In the following, we show that in the zebrafish visual error signals are not necessary for the early development of the neural integrator, and present a model for how the integrator can develop using a feedback loop through the cerebellum. Integrator output is low-pass filtered at the cerebellum through an internal model of the oculomotor plant before being fed back. This cerebellar feedback then serves as a teaching signal at the VPNI by virtue of segregation of local and global feedback signals onto separate neuronal compartments. In this manner we show how an initially randomly connected forgetful network can learn the appropriate recurrent connection weights necessary to perform temporal integration. As the feedback strictly uses internally generated signals, it achieves a form of bootstrap learning.

## 2 Results

We first present the experimental finding that visual error signals do not play a significant role in the early development of the zebrafish VPNI. We then lay out a new hypothesis for how this self-sustained development could occur through a low-pass filtered feedback of the aggregate integrator activity via bootstrap learning. We begin presenting this hypothesis with a simple model that captures the essence of our proposal. Next, we present two real-time stabilization mechanisms that complement the synaptic tuning mechanism and help generate robust temporal integration. We then demonstrate how the tuning mechanism is intimately related to a segregation of local and global feedback inputs onto separate neuronal compartments, and show that these mechanisms function appropriately in a spiking network composed of conductance-based neurons. Finally, we incorporate all of the proposed components into a model of the oculomotor system that includes the VPNI, the cerebellum, and motoneurons to demonstrate how realistic tuning curves and response dynamics can arise through our bootstrap learning hypothesis.

### Early development of the neural integrator is independent of visual inputs

Given prior work suggesting visual error signals are important for integrator plasticity, at least in adult animals, we first asked if visual feedback was important for the early development of the neural integrator. We compared the integration time constant of light-reared animals to those of dark-reared animals. Light-reared animals were raised under standard conditions with a 12 hour light-dark cycle. Sibling dark-reared animals were raised in similar conditions but only exposed to ambient light for a minute or two during preparation for eye tracking. When measured at 5 days post fertilization, within 24 hours of the development of saccade eye movements, animals in both groups made spontaneous saccades in the dark, with saccades occurring every 30.1 ±11.6 seconds for light-reared animals, and 35.9 ±14.1 seconds for dark-reared animals (Fig. 1A). After saccades, the eyes generally exhibited a relatively rapid slide to a period of fixation with moderate to minimal drift towards baseline, a drift pattern that reflects the presence of a leaky integrator (Fig. 1A).

**Figure 1:**
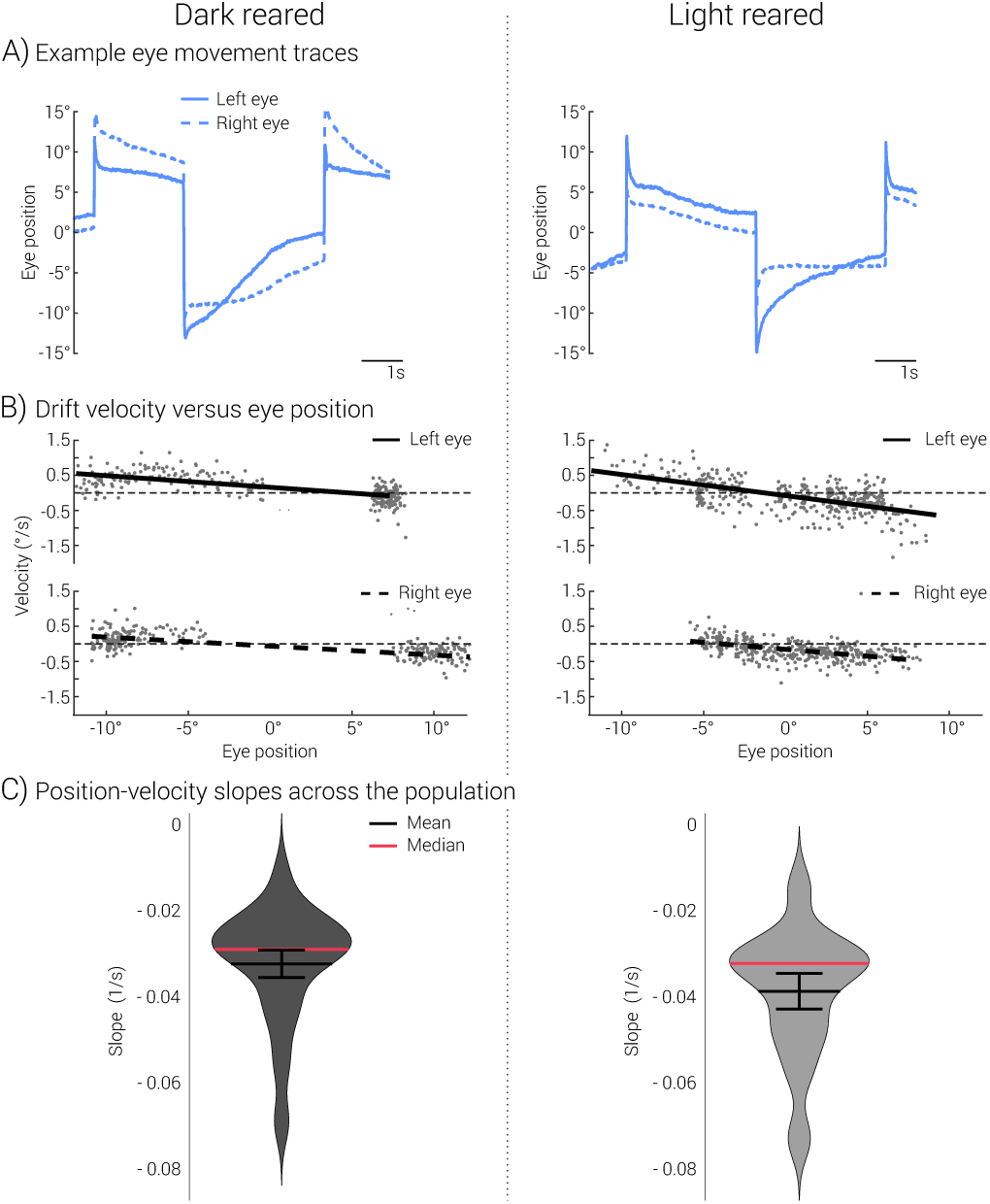
Learning to integrate is independent of external feedback. **A)** Typical spontaneous saccades and fixations of zebrafish larvae reared in the dark (left column) or reared on a standard light-dark cycle (right column). **B)** Example plots of fixation drift velocity versus fixation position for both eyes of a dark-reared (left) and light-reared (right) animal. **C)** Violin plots of drift dynamics from dark-reared and light-reared animals.

Quantification of this drift revealed that there was little difference in integrator functionality between the two populations. To quantify this drift pattern, we plotted drift velocity as a function of eye position and identified the inverse slope of the best linear fit as an integration time constant (Fig. 1B). The analysis revealed that light-reared animals had comparable integration time constants to dark-reared animals (Fig. 1C, *p* = 0.22; 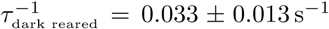, *n* = 18; 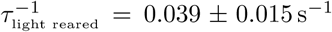, *n* = 13. This demonstrates that the initial tuning of the neural integrator must use a mechanism distinct from the retinal-slip mediated plasticity shown to be important for oculomotor plasticity in the adult setting.

### Conceptual model for bootstrap integrator learning

Motivated by the above finding, we propose a boot-strap learning mechanism that would allow for the development of temporal integration without the use of external error signals. To illustrate the learning paradigm we consider first a simple model with two neurons, one representing an integrator population, and the other a teaching population (Fig. 2A). In the example of the zebrafish, the integrator population is identified by the VPNI, and the teaching population is identified as the cerebellum. Both populations receive the same stimulus (*r*^S^) driving saccadic eye movements. The cerebellum receives excitatory input from the integrator, and the integrator receives a teaching signal from the cerebellum with mixed excitatory and inhibitory synaptic pathways (through vestibular neurons, not shown). The integrator is recurrently connected to itself through an excitatory connection.

**Figure 2:**
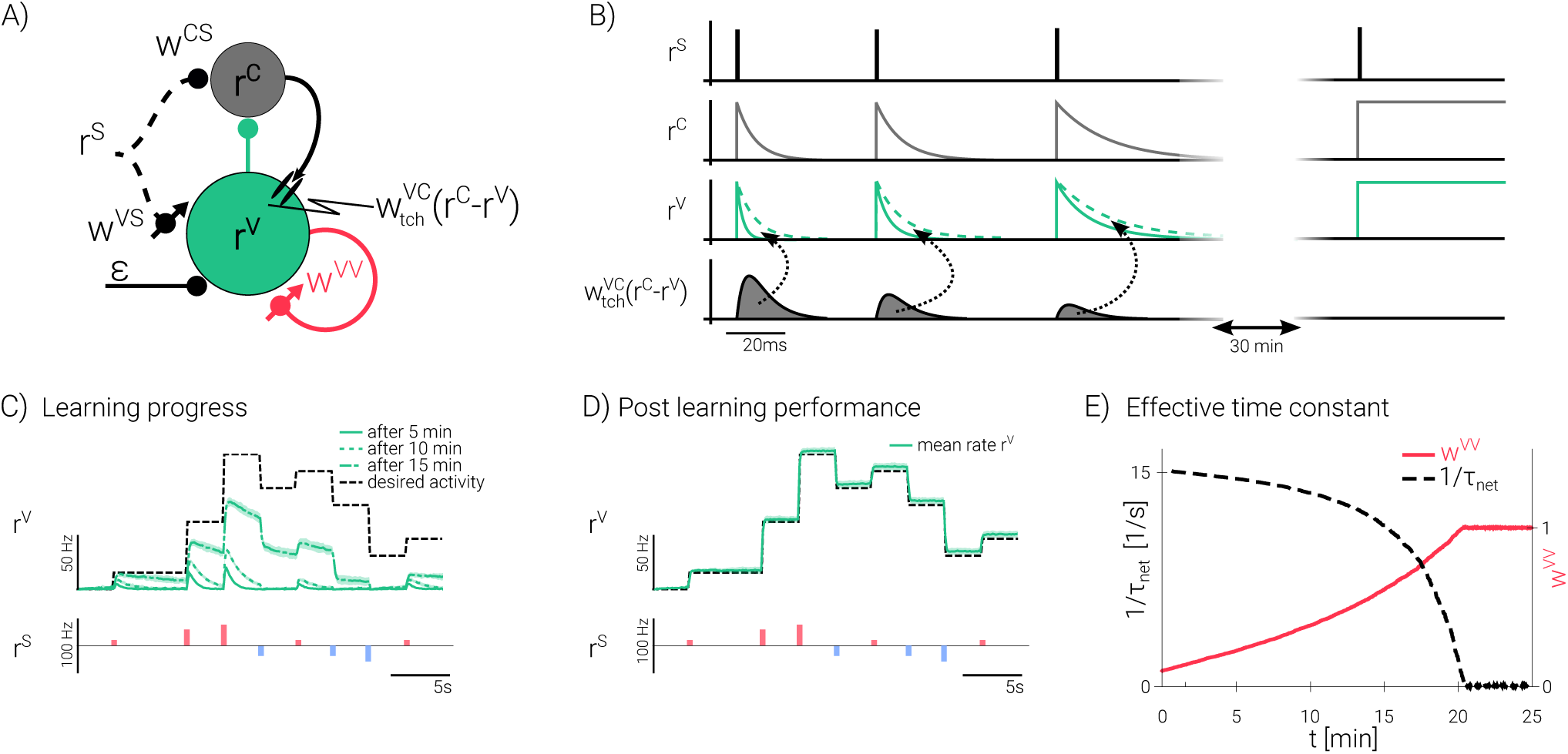
Integrator learning with a delayed feedback acting as target activity. **A)** Bootstrap learning in an integrator population (green) is based on a delayed feedback signal from a teaching population (gray). The learning is based on a delayed feedback signal r^C^ that represents a low-pass filtered version of the integrator activity r^V^. Both r^C^ and r^V^ are driven by saccade inputs r^S^ to be integrated in time (via synaptic strengths w^CS^ and w^VS^, set in Eqs 1 & 2 to 1 for the sake of simplicity, but see Methods for also learning these synapses). To keep its value, r^V^ feeds back onto itself through the weight w^VV^ that is learned by an error-correcting plasticity rule (Eq. 3). **B)** The feedback r^C^ transforms at the level of the postsynaptic voltage to an error signal 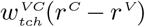 that pushes r^V^ towards r^C^. After 30 min saccade stimulations the decay was eliminated and r^v^ = r^C^ = ∫ r^S^ (rectangles). **C, D)** Averaged traces of r^V^ (top, green, averaged across 20 traces) with positive (bottom, red) or negative (blue) saccade pulses r^S^, after early learning (C; 5, 10, 15 min) and late learning (D; 25 min, simulations of Eqs 7–9 that include noise; black dashed: true integral of the saccade commands, i.e. right-hand side of Eq. 4). **E)** During learning, w^VV^ converges to 1 (solid red) while the network integration time constant becomes infinite (1/τ_net_ = 0, dashed, Eq. S2). The learning rate is chosen to get stable learning with a 0.5 Hz saccade rate, but slower learning is also possible.

The performance of the integrator is critically dependent on the strength of the local excitatory recurrent feedback *w*^VV^. When mature, the integrator population feeds back onto itself with sufficient strength to integrate the saccadic input *r*^S^ and sustain its own activity (*r*^V^) during the subsequent fixation. Perfect integration occurs if the synaptic feedback strength is one, *w*^VV^ = 1. Initially, however, the feedback strength may be mistuned; if it is too small, for instance, the integrator is forgetful (Fig. 2B).

Learning in this paradigm is driven by the difference between the activity of the integrator and the cerebellum. We posit that the cerebellum low-pass filters the input from the integrator; hence, if the integrator is forgetful, during the attempted fixation the cerebellar signal (*r*^C^) will always decay more slowly than the integrator signal (*r*^V^) so that we have *r*^C^ > *r*^V^. To learn how to integrate, then, we determine at the integrator population the difference between the delayed feedback signal from the cerebellum and integrator activity (*r*^C^−*r*^V^), scale it by a teaching weight 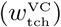, and use this difference signal to drive plasticity of the local recurrent feedback strength *w*^VV^. The strength *w*^VV^ is modified until the control and integrator activity become equal; i.e., when perfect integration is obtained (Figure 2B, right). Our self-supervised learning scheme effectively allows the integrator to correct its forgetfulness in a manner reminiscent to pulling oneself up by their own bootstrap. We therefore refer to this self-supervised scheme as bootstrap learning.

More formally, the dynamics for integrator and control population can be written as (setting neuronal time constants and other adaptable weights to 1 for simplicity; see Methods)

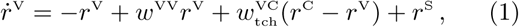

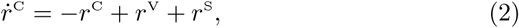

where both populations receive the brief driving stimulus *r*^S^ (Fig. 2A). Here, 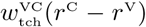 is the teaching current that contributes to the stabilization both on the fast neuronal and the slow synaptic time scale as explained below. The plasticity is postulated to follow the error-minimizing gradient rule (Methods)

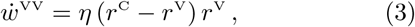

with *η* being the learning rate. Learning only stops when plasticity drives the activities to become equal, *r*^V^ = *r*^C^, and hence 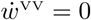. But equality also implies 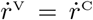, and for Eqs 1 and 2 to generally hold one infers that *w*^VV^ = 1. In this case the activity dynamics reduces to 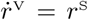 and 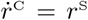, and integrating the first equation yields

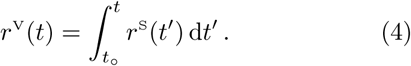

Simulating a noisy version of Eqs 1–3 (Methods) leads to the development of a stable integrator with a network time constant that in fact grows to infinity (Fig. 2C-E). In the full zebrafish oculomotor model below, we show that the bootstrap procedure can also be nested to simultaneously learn the low-pass filtering *r*^C^(*t*) in the cerebellum based on noisy and distributed VPNI activity.

### The delayed feedback signal also provides real-time stabilization

Proper integration over a wide dynamic range requires a precision of tuning that might be challenging to achieve and maintain for biological systems. Various robustness mechanisms have been proposed to deal with this fine tuning problem, including synapto-dendritic nonlinearities and (Koulakov, Raghavachari, et al. 2002) and delayed negative feedback (Lim and Goldman 2013). We show in the section and the next that our learning mechanism can work in concert with these robustness mechanisms.

The synaptic teaching signal introduced above drives learning on a multiple minutes time scale, but also provides a moment-to-moment delayed derivative feedback that helps to alleviate the fine-tuning concern. On the one hand, the integrator activity *r*^V^ is directly fed back via plastic synaptic strength *w*^VV^ that is modified to sustain the activity in the absence of input. On the other hand, it is indirectly fed back via lowpass filtered activity *r*^C^ and fixed connection strength 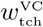 to form a negative error feedback 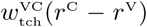, see Fig. 2A and Eq. 1. At any given moment, this difference signal dampens deviations in integrator activity from its low-pass filtered version: whenever *r*^V^ deviates from *r*^C^, the signal 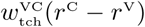 counteracts on a fast time-scale. Whenever the recurrent feedback strength *w*^VV^ is mistuned and, as a consequence, the activity diffuses away, the difference signal counteracts (Fig. 3A-C). The effect of this correction is visualized by varying 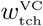: the weaker the strength of this negative derivative feedback, the stronger the drift.

**Figure 3:**
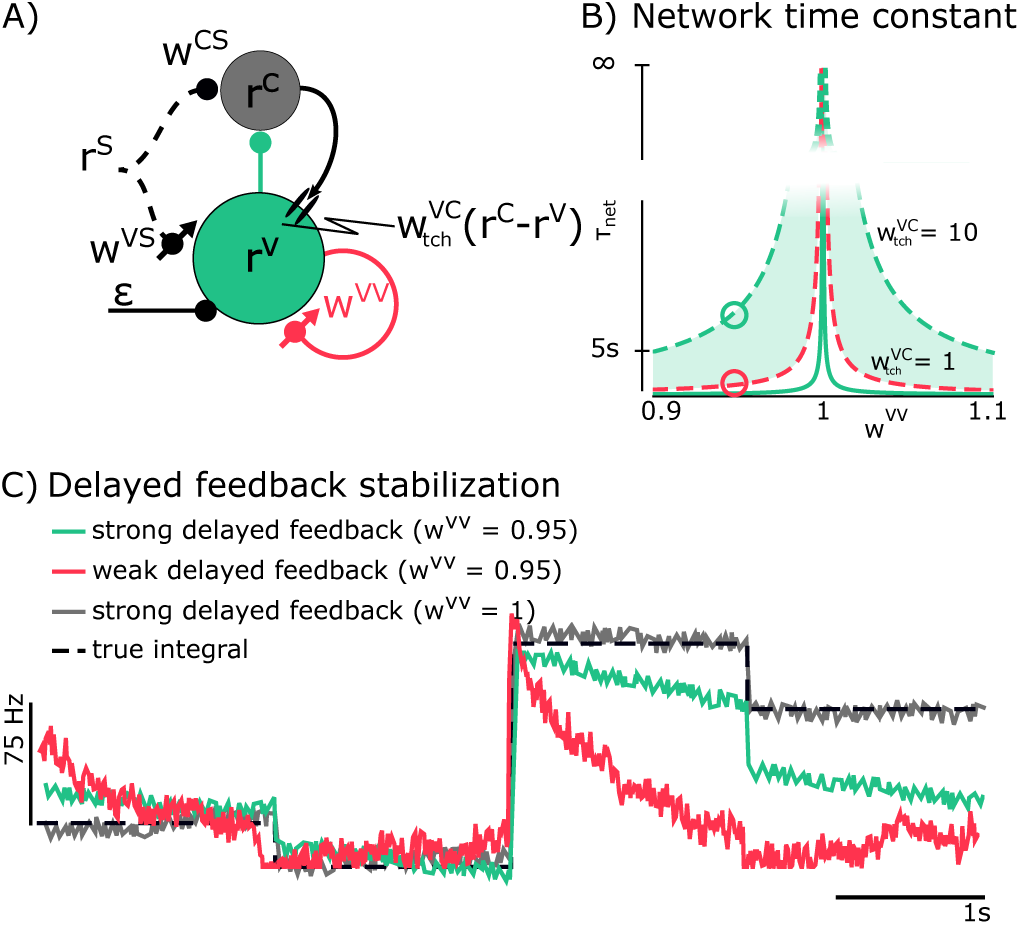
The delayed feedback signal 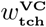 also stabilizes integration with respect to mistuned feedback weights (*w*^VV^) and with respect to noise (*ε*). **A)** The feedback r^C^ that at the level of postsynaptic voltages is expressed as an error signal 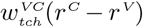, also stabilizes the network integration in the presence of noisy input (ε, black arrow). **B)** Network time constant τ_net_ of the coupled system (Eq. S2) as a function of the recurrent synaptic strength w^VV^ and different teaching strengths 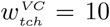 (dashed red), 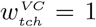 (dashed green) and 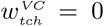 (solid green). For w^VV^ = 1 the network is a lossless integrator. **C)** Example traces of the integrator activity r^V^ with mistuned recurrent feedback (w^VV^ = 0.95) and different teaching strengths (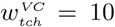green;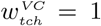, red; corresponding to the circles in B). After perfect learning, when w^VV^ = 1, the integrator activity follows the true integral (black dashed) and is also robust against weak noise (grey, 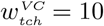).

While the stabilization acts on the fast time scale of seconds, the teaching signal simultaneously serves as a drive for synaptic plasticity that adjusts the feedback strength towards *w*^VV^ = 1, as explained above. As we show later, the self-supervised learning of the integrator property can naturally be interpreted in a two-compartment neuron model where synapses on the dendritic tree learn to predict the somatic firing (Urbanczik and Senn 2014). Before describing this dendritic implementation, we consider populations of principal and interneurons and show that a nonlinear recruitment of the population neurons further stabilizes the persistent activity.

### Learning in a population and stabilization through nonlinearities in recurrent feedback

In the above simplified model we considered a single integrator unit that in biology would be represented by a population of neurons, typically composed of excitatory and inhibitory neurons. Learning in the presence of multi-synaptic feedback loops in a population of neurons is more challenging and we show that our boot-strap learning can also deal with this situation (Fig. 4A). Additionally, we incorporate nonlinear synaptic interactions within the Integrator such that sequential recruitment of neurons provides an additional stabilization mechanism (Seung et al. 2000; Brody, Romo, and Kepecs 2003).

**Figure 4:**
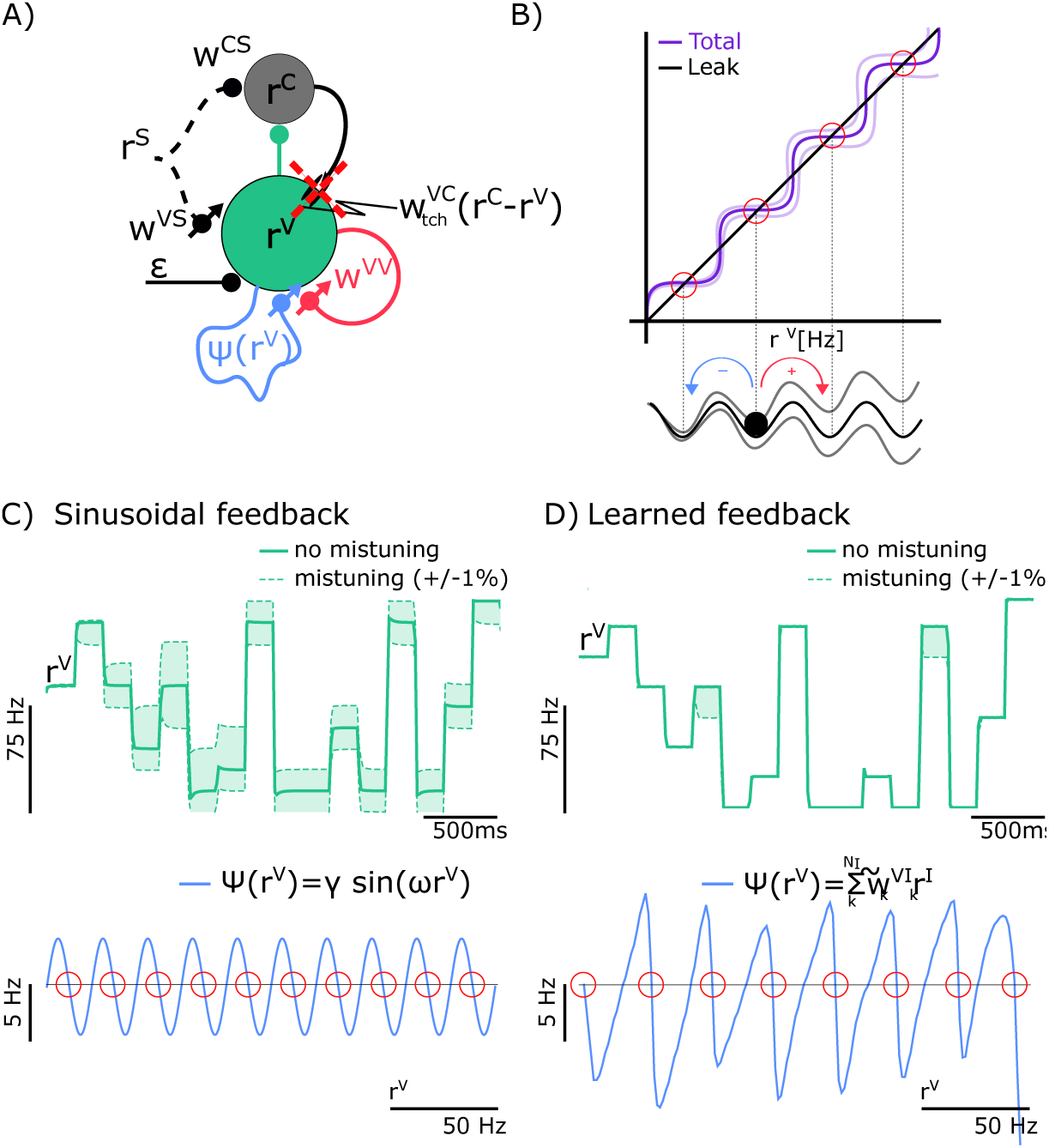
Formation of stabilizing nonlinear feedback through recruitment. **A**) With a nonlinear feedback learned by recurrently recruiting interneurons endowed with synaptic short-term depression (wiggled blue line, ψ(r^V^), see Eq. 6), the integrator remains stable even when the teaching signal is cut after learning (red cross). **B)** For stability, the net recurrent feedback, w^VV^r^V^ + ψ(r^V^) (purple lines with different values for w^VV^), needs to cross the identity line r^V^ (black) from above (red circles). **C)** Example for ψ (blue sinusoidal function of r^V^) yielding 10 stable fixed points between 0 and r^max^ (red circles). **D)** If ψ(r^V^) dynamically develops due to the plasticity of the depressing synapses from inter-to-principal neurons (w^VI^), stability even improves (top, simulation of Eqs 8, 10 and 13–16).

In the population model of the Integrator, we consider a system with integrator principal neurons that provide feedback connections within the integrator and interact with other circuits, and integrator interneurons that only receive and provide input to the principal neurons (Fig. 4A). In the following, we first describe for illustrative purposes how learning can be achieved with an idealized form of nonlinear feedback through the interneuron population, and we then show how the appropriate structure of inter- and principal neurons can evolve through learning.

Through the recruitment of neurons in a population an additional stabilization can be achieved, by effectively finding a population input-output transfer function that is ‘wiggling’ around the identity. To explore the idea, we split the population feedback into a linear, direct feedback of strength *w*^VV^*r*^V^ between principal neurons, plus a nonlinear indirect feedback of strength *ψ*(*r*^V^) through interneurons. The integrator dynamics then reads as

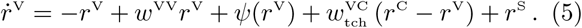

To study the benefits of local nonlinearities, we first consider a sinusoidal modulation (*ψ* = sin). Learning then leads to an overall feedback close to unity, *w*^VV^*r*^V^ + *ψ*(*r*^V^) ≈*r*^V^, with the direct feedback strength *w*^VV^≈1 as before. But now, for rates 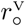 with 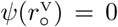 and negative slope 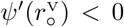, stable attractors develop (Fig. 4B).

We next describe how the interneuron population can be organized to generate the nonlinear feed-back described above. In this simplified example, we considered two integrator principal neurons with average rate *r*^V^ that drive a population of *N*_I_ interneurons. Both principal and interneurons fire with rates determined by a threshold-linear transfer function, 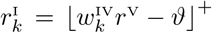, as observed for VPNI neurons (Fig. 5A, see also Aksay, Baker, et al.). The principle-to-interneuron weights 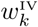 are sampled from a heavy-tail distribution, assuring that the number of recruited interneurons linearly increases with *r*^V^ (Fig. 5C and Methods, Eq. 17). The interneurons project to the principal neurons through either excitatory or inhibitory weights 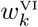 that undergo short-term synaptic depression. The synaptic release rate of these interneurons is a saturating function of the presynaptic rate *r*^V^ (Fig. 5C). With this, the indirect feedback now becomes

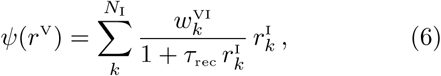

where *τ*_rec_ determines the recovery time from the synaptic depression, *N*_I_ the number of interneurons recruited at VPNI rate *r*^V^, and 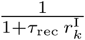 is the depression factor that down-scales the absolute synaptic weight 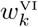 from interneuron *k* to the principal neurons (neglecting the ‘use’-parameter, see Methods, Eq. 14, and Tsodyks and Markram).

**Figure 5:**
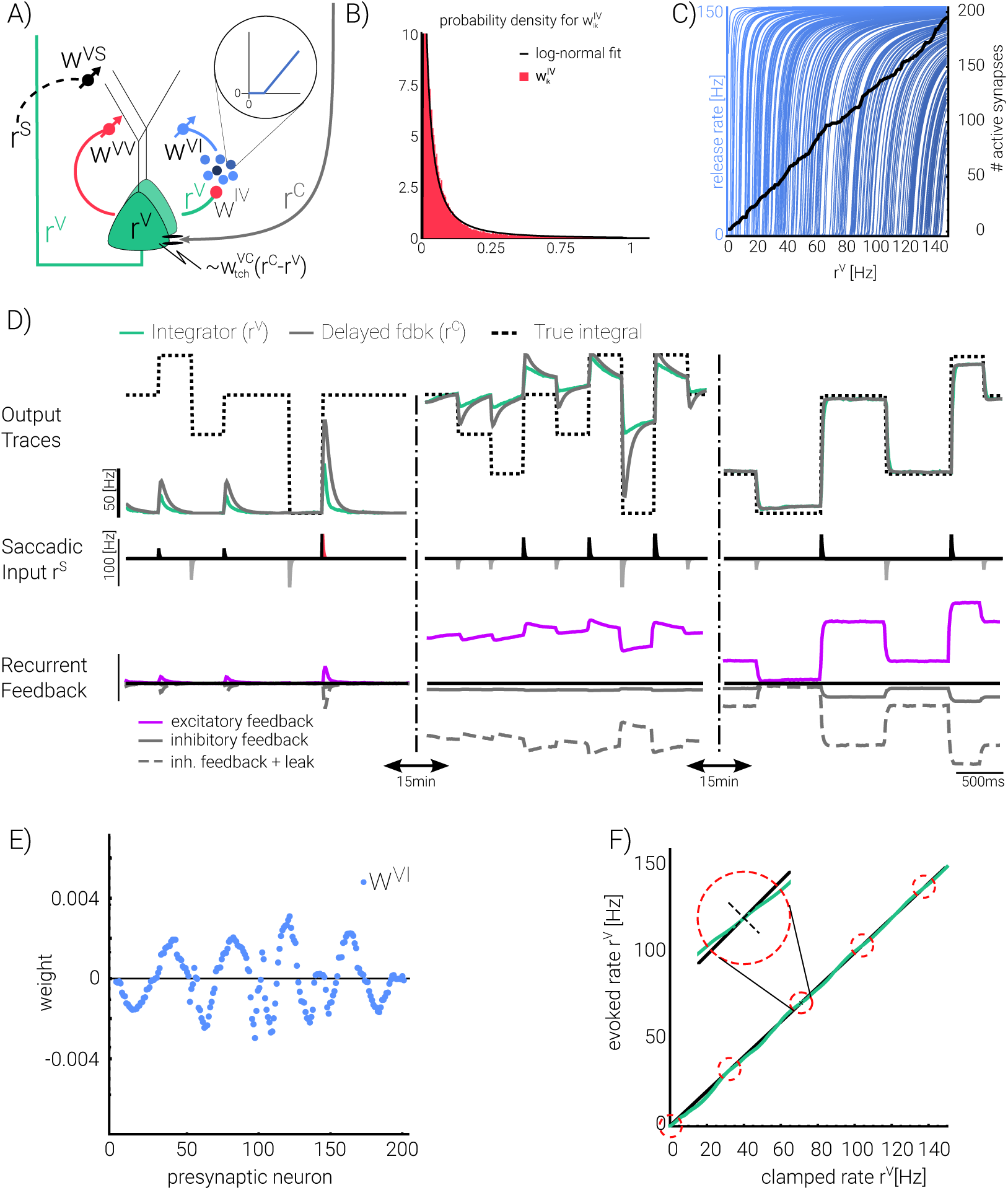
Two-compartment neuron implementation and emergence of balanced states. **A)** Conductance-based somatic input naturally implements the delayed teaching signal 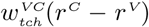 from Eq. 5 as a somatic teaching current. Recurrently recruited interneurons (blue, with identical current-to-rate transfer functions, inset) project to the dendritic compartment of the principal neurons through synapses with short- and long-term plasticity. **B)** The distribution of the principal-to-interneuron weights, 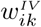 (red), that lead to a linear recruitment of the interneurons is roughly log-normal (black). **C)** Contribution of the N_I_ = 200 interneurons as a function of the population rate 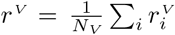 with N_V_ = 2 principal neurons (for a more realistic number/VPNI structure see Fig. 6/7). Blue lines represent the individual terms in Eq. 6, interpreted as synaptic release rate (cf. Eqs 13-15). The number of recruited interneurons is roughly linear in r^V^ (black). **D)** Evolution of r^V^ (green) and r^C^ (black) in response to positive and negative saccade inputs (middle traces). Lowest traces: Repacking of the direct 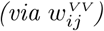 and indirect 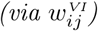 feedback currents into groups excitatory (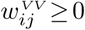 and 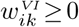, purple) and inhibitory (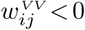 and 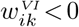, grey) currents. After learning (rightmost), the total excitatory current is mirrored in the inhibitory current and is fully balanced by the sum of inhibitory and leak current during the stabilization period (dashed grey, see also Supp. Info. S1.5). **E)** The weights 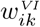 from the inter-to-principal neurons evolved during learning such that they stabilize the firing rates in the absence of inputs. **F)** Open-loop mean firing rate r^V^ that would be evoked in the population of principle neurons if these are clamped to a given value (green curve: w^VV^r^V^ + ψ(r^V^) as a function of r^V^). Downwards crossings with the diagonal (black) yield stable fixed points for r^V^ (inset, cf. Fig. 4).

The absolute inter-to-principle neuron strengths 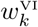 are plastic with the same error-correcting learning rule as for the previously described direct feedback weights, 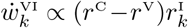. With these additional nonlinearities introduced by the interneuron recruitment and the synaptic depression, the bootstrap learning can still form a recurrently connected neuronal population that faithfully integrates saccade inputs (Fig. 4D). Indeed, the learning mechanism is able to find weights 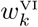 to yield a feedback loop through the interneuron population that is qualitatively similar to the idealized sinusoidal form introduced before. The number of attractors is limited by the number of neurons in the VPNI population and it also depends on the distribution of saccade inputs *r*^S^ used during learning (Supp. Fig. S1).

### Integrator learning with two-compartment spiking neurons

Before moving to the full model, we explain how the teaching current 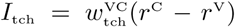 in Eq. 5, i.e. the deviation of the integrator activity from its delayed feedback via cerebellum, is generated biophysically and how it can serve as a drive for plasticity. For this we endow our VPNI integrator neurons with a somatic and dendritic compartment. The direct VPNI feedback with connection strength *w*^VV^ projects to the dendritic compartment, as does the indirect feedback through the interneurons and the saccade input with corresponding connection strengths. The cerebellar activity *r*^C^ projects through conductance-based direct inhibition and indirect excitation to the somatic compartment (Fig. 5A).

The key property of the membrane biophysics is that the total conductance-based cerebellar input, *I*_tch_, defines a reversal potential (called the matching potential *U*_M_) where excitatory and inhibitory currents exactly balance out so that the total cerebellar input current vanishes (Urbanczik and Senn 2014). As a consequence, if the recurrent input to the VPNI neuron already imposes the same postsynaptic voltage in the soma, *U* = *U*_M_, the cerebellar input vanishes, *I*_tch_ = 0, and the somatic voltage stays at its value. Formally, the total cerebellar input current is proportional to the difference between the matching potential and the so-matic voltage, 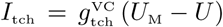, with proportionality factor being the sum of the excitatory and inhibitory conductances 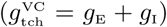 induced by the cerebellum.

The biophysics of the cerebellar input is reflected in the form of the teaching current in the simplified model of Eq. 5, 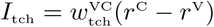. The matching potential *U*_M_ is induced by the cerebellar activity *r*^C^, and the postsynaptic voltage *U* is, in the absence of saccade commands, driven by the VPNI activity *r*^V^. Importantly, the VPNI and cerebellar activities are identical, *r*^V^ = *r*^C^, when *U* = *U*_M_, and hence the teaching current in the simplified model faithfully represents the biophysics of the cerebellar input. Learning is driven by the difference between the firing rate established by cerebellar inputs minus the firing rate caused by the dendritic input, the so-called dendritic prediction error, that becomes a fixed fraction of the difference rule 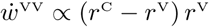 as expressed in Eq. 3 (see also *r*^C^ − *r*^V^. This leads to the error-correcting learning Methods, Eqs 23–28).

To demonstrate the functionality of this framework, we simulated a VPNI network composed of two-compartment principal neurons and a large population of VPNI interneurons that project through depressing but plastic synapses back to the principal neurons. After 30 minutes of biological simulation time a robust integrator evolved (Fig. 5D), with the well-known balancing of excitation and inhibition (Denève and Machens 2016). The recurrent excitation is balanced by the overall recurrent inhibition including the leak, as required by the very nature of the sustained activity between saccades, formally *w*^VV^*r*^V^ + *ψ*(*r*^V^) − *r*^V^ = 0 in Eq. 5. The learning rule can also cope with the population of interneurons by appropriately adapting the inter-to-principal connections, 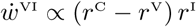, such that the overall feedback exploits the nonlinear recruitment and robustly stabilizes the delay activity as explained above (Fig. 5E,F).

The integrator also robustly emerged when both the principal and interneurons were spiking with an instantaneous Poisson rate 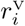 and 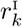, respectively (Fig. 6). Here, to reduce fluctuations, the number of principal neurons was increased to a more realistic 100. Learning was driven by the spiking version of the dendritic predictive plasticity rule for all the spiking inputs from the principal neurons themselves, the interneurons, and the saccadic units (Eq. 28 and Supp. Info. S1.5). During the ongoing random saccade stimulations, all these synaptic weights jointly learn to match the spike-induced postsynaptic potential of a principal neuron to the cerebellum-imposed matching potential in real time (Fig. 6B).

**Figure 6:**
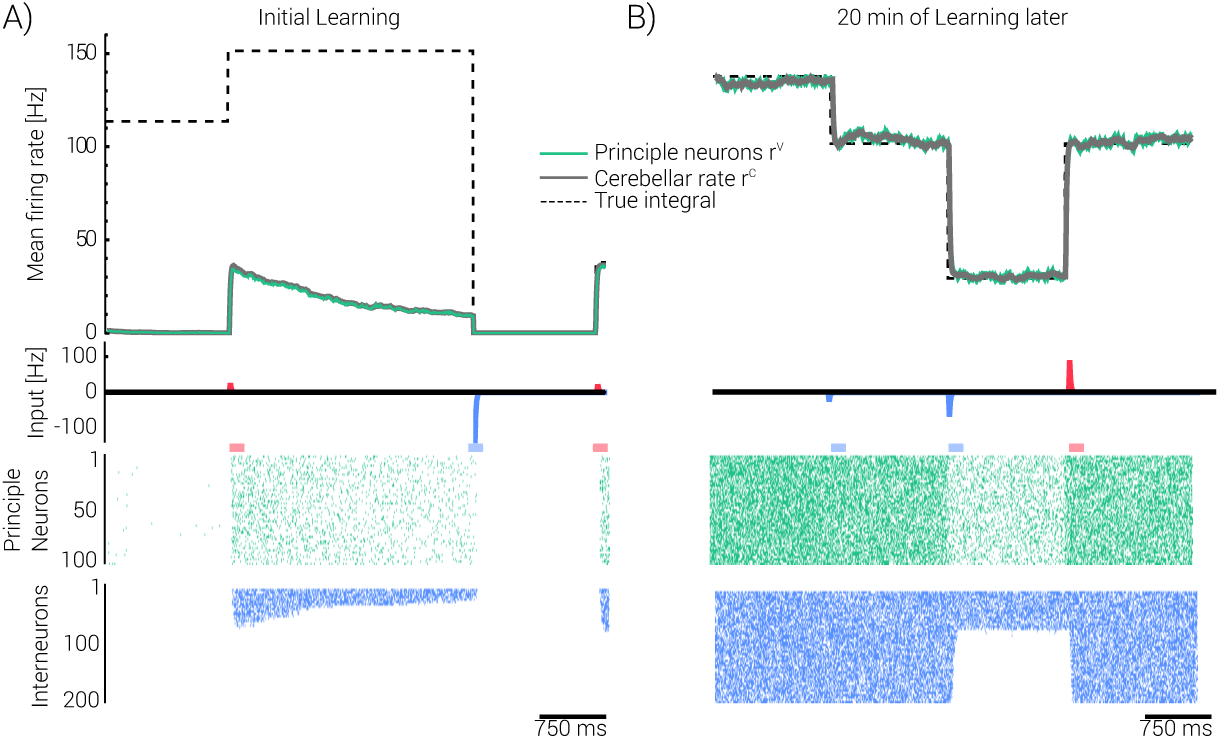
Stable integrator learning with spiking two-compartment neurons. Same simulations as in Fig. 5D, but with spikes triggered by the instantaneous Poisson rates for principle and interneurons, 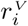 and 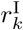 (Eq. 22). Lower panels: spike raster plot of a fraction of principal neurons (green, N_V_ = 100) and release raster plot of the interneurons (blue, N_I_ = 200). Dashed line: ∫dt r^S^(t). **A)** Initially, the population firing rate 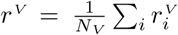 (green solid: mean; green dashed: ±standard deviation) deviates from the cerebellar teaching signal r^C^ (solid grey). **B)** After 20 min of exposure to the saccade inputs (r^S^, middle trace) the VPNI principal neurons learned to integrate.

### Learning in the cerebello-integrator circuit of the larval zebrafish

We next applied the idea of bootstrap learning to a more realistic model of oculomotor temporal integration in the developing zebrafish. In the above, we first showed that the oculomotor neural integrator is able to tune up in the absence of visual inputs (Fig. 1), presented a conceptualization of how this could be accomplished through a delayed internal feedback that provides a teaching signal (Fig. 2), showed how this delayed feedback could by itself (Fig. 3), and when coupled with local feedback nonlinearities (Fig. 4), be used also to provide real-time network stabilization. We further demonstrated how at the cellular level the stabilizing plasticity rule emerges from the distribution of the cerebellar teaching feedback and the local VPNI feedback across somatic and dendritic compartments of neurons within a population (Figs 5 and 6). Extension of these concepts to a more realistic setting of the zebrafish, however, requires that we tackle two additional challenges beyond the generation of persistent firing: namely, the development of the appropriate neuronal tuning curves and the generation of the appropriate delayed feedback signal. Specifically, the first challenge is to have an integrator network also learn the slope-threshold organization apparent experimentally over a bilateral population (Aksay et al, 2000); the second challenge is to compile from the threshold-linear responses of developing integrator neurons a teaching signal that evolves to represent the desired eye position. Furthermore, the tuning curves, teaching signals, and persistent firing need to mature in tandem.

The key to dealing with these challenges is to allow cooperative development of the different brain centers involved in producing oculomotor responses. The full model consists of four neural populations (Fig. 7A): the integrator (VPNI), the saccade generator (Sacc), the cerebellum (CE), and the abducens motoneurons (AN). The VPNI consists of an ipsi- and contra-lateral population of VPNI neurons. Initially, the integrator neurons are randomly connected among themselves, and they randomly project to motor neurons in the abducens, but these neurons receive structured teaching feedback from the cerebellum. To develop properly, the system must simultaneously find the appropriate weights of modifiable synapses at several key locations: at the synapses from the saccade generator onto each of the other populations, and at the synapses from the integrator onto both the abducens and the cerebellum. The teaching signals that regulate the simultaneous adaptation at the disparate synaptic sites all arise from the cerebellum (Fig. 7A): the cerebellar projection 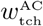 regulates adaptation of synapses from the saccade generator and integrator onto the abducens; the cerebellar projection 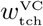 regulates plasticity of synapses from the saccade generator onto the integrator and of synapses between integrator cells; and the cerebellar self-projection 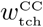 regulates plasticity of the synapses from the saccade generator and integrator onto the cerebellum (Eqs 38, 33 and 29). Note that these teaching signals themselves evolve over time as the inputs to the cerebellum from the saccade generator and integrator are modified.

**Figure 7:**
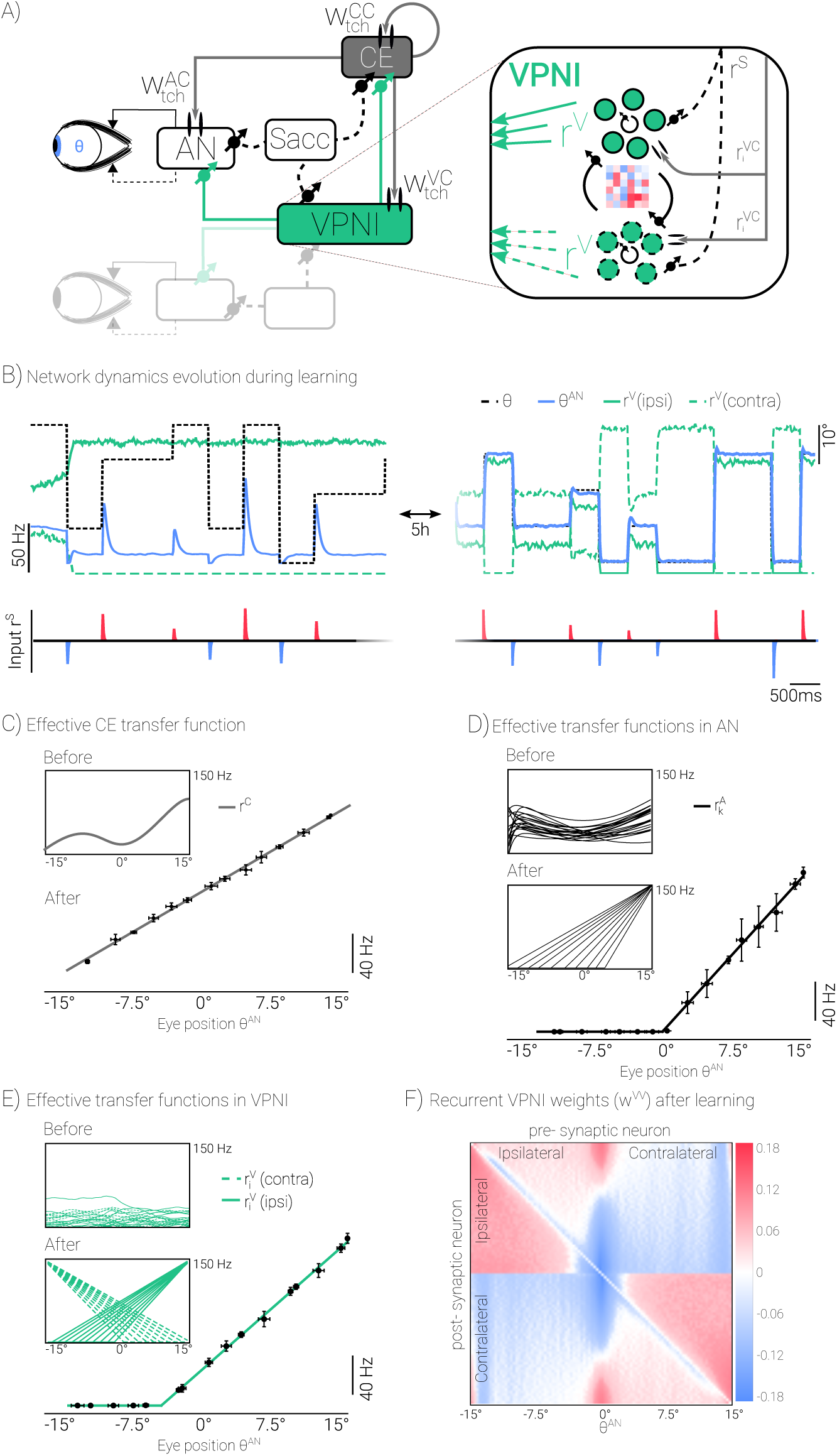
Self-supervised learning in CE, VPNI and AN produces stable integration with observed connectivity and transfer functions. **A)** Full circuit of the cerebellum (CE), the abducens nucleus (AN) and the VPNI. All synapses marked by a crossing arrow are plastic. The evolving CE provides teaching signals (weighted by 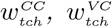 and 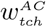) to itself as well as to the VPNI and AN neurons. Inset: Ipsi- and contralateral VPNI neurons that are recurrently connected, with initially random connectivity matrix (color code). They receive saccadic input r^S^ and a teaching signal r^VC^ from the CE. **B)** 5s example traces of the saccade integral (black dashed), the produced eye position (blue) and the firing rate of an ipsi (green) and contralateral (green dashed) VPNI neuron, before (left) and after 5h of stochastic saccade stimulation (right). Below: Corresponding saccade inputs r^S^. **C)** Cerebellar position-to-rate transfer function before (inset) and after learning, with error bars for the produced eye positions and rates. **D-E)** Example of a AN (D) and VPNI (E) position-to-rate transfer function after learning with error bars for the produced eye positions and rates. Insets: Transfer functions for all AN and VPNI neurons before and after learning. **F)** Recurrent weights within the VPNI developed ipsilateral excitation and contralateral inhibition, plotted against the eye position θ at which the pre- and postsynaptic neurons start firing.

With the specific choices for the above parametrization of the cerebellar teaching signals we imposed a minimal set of biologically-inspired constraints upon the model. A first set of constraints could originate from proprioceptive inputs from the oculomotor muscles that sets the appropriate internal model in the cerebellum (Fiorentini and Maffei 1977; Steinbach 1987). We posited that proprioceptive inputs help generate an internal model of the aggregate response in the abducens and plant by setting the filtering properties of the cerebellar self-projection loop (Eq. 29). We also posited that proprioceptive inputs help calibrate the range of abducens firing rates appropriate for covering the oculomotor range, and we assumed that the muscle force, being the weighted sum of the abducens firing rates, is matched to the oculomotor range and the muscle properties (Eq. 40). A second set of constraints could originate from genetic factors. One of these were that the inputs from the cerebellum were excitatory (through interneurons) to one half of the integrator population and inhibitory to the other half of the integrator population through a monotonically increasing distribution of ‘teaching weights’ (Eq. 33). Another constraint is that the cerebellar input similarly ‘teaches’ the abducens neurons again through monotonically increasing distribution of teaching weights (Eq. 38).

To test whether this full model can cooperatively develop a stable integrator with the appropriate tuning curves, we started out with a system with initially mistuned random weights, and simultaneously ran the neuronal dynamics (Eqs 29, 31 and 36) and the synaptic plasticity rules (Eqs 30, 34 and 39) during a sequence of saccades. To relate to the experiments, we further plotted the cerebellar, AN and VPNI activities as a function of the produced eye position *θ*^AN^. Due to the random initialization, the teaching signals from the cerebellum to itself, to the VPNI and the AN, were initially inappropriate and the cerebellar activity, which should eventually represent internal model of the eye position, did not reflect the desired eye position, *r*^C^(*t*) ≠ *θ*(*t*) where 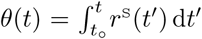. Furthermore, the true eye position produced by the abducens neurons (Methods) was incorrect, (Fig. 7B), and the VPNI and AN position-to-rate transfer functions were distorted (Fig. 7C insets ‘Before’).

With ongoing saccadic stimulation and plasticity at the various synapses, however, the network learned to properly integrate, and the produced and desired eye positions eventually matched across the full range, *θ*^AN^ ≈ *θ* (Fig. 7B). In doing so, the position-to-rate transfer function of the CE became linear, consistent with the idea that the CE represents an internal model of how the abducens and plant transforms input signals to eye position (Fig. 7C). The position-to-rate transfer functions of the AN neurons, 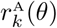, and VPNI neurons, 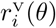, developed a positive threshold slope correlation (‘After’ in Fig. 7D,E) as observed in the real preparation (Pastor, Torres, et al. 1991; Aksay, Baker, et al. 2000).

Simultaneously, a highly structured VPNI connectivity matrix developed out of the originally random connection strengths (Fig. 7F and Supp. Fig. S6). Generally, VPNI neurons became strongly excited by ipsilateral VPNI that had a lower recruitment threshold as a function of *θ*^AN^, and became weakly excited or even inhibited by ipsilateral VPNI neurons with a higher recruitment threshold. Inputs from contralateral VPNI neurons, on the other hand, typically were inhibitory. Together, the pattern that emerged from the bootstrap learning procedure largely reflected the push-pull organization of the oculomotor integrator evidenced by earlier work (Fisher et al. 2013).

## 3 Discussion

We have experimentally shown that the eye position integrator in zebrafish larvae develop equally well in dark-reared animals without any visual feedback and in particular without exploiting retinal slip (Murakami and Cavanagh 1998). We explained this phenomenon by a general model that autonomously learns to temporally integrate. For the example of the oculomotor system in the zebrafish we showed how the cerebellum may act as an intrinsic feedback control for learning the integrator dynamics and for adapting the motor commands.

### Learning based on dendritic predictions

We have proposed a model for integrator learning that takes advantage of the spatial extent of integrator neurons. Previous models of integrator learning, and the great majority of learning models in general, assume single-compartment neurons for convenience and tractability. However, such models ignore the very real possibility that the signals driving learning and the signals being modified arrive at or are present on different parts of the neuron, be it proximal vs distal dendrites, or different dendritic branches (Urbanczik and Senn 2014). We argue that this heterogeneity should be viewed as a feature that can be exploited for learning, and propose a scheme wherein cerebellar teaching signals arriving at or near the soma can drive the plasticity of recurrent integrator signals arriving at more distal dendritic regions. Potential heterogeneity in inputs is hinted at in a recent ultrastructural study of identified integrator neurons, which suggested differing spatial distributions for local feedback from integrator neurons vs global input from other sites (Vishwanathan et al. 2017). Future studies may consider the precise distribution of cerebellar inputs (through vestibular nuclei) onto integrator neurons, and the potential for independence in the plasticity of different input streams via segregation of those streams onto differing dendritic locations.

### Cerebellar control

We propose that the cerebellum generates the teaching signal necessary to drive plasticity at the level of the integrator and the abducens. Such a role seems plausible given the involvement of the cerebellum in a wide range of oculomotor learning behaviors, the presence of connections from integrator neurons to the cerebellum (Lee, Arrenberg, and Aksay 2015; Kolkman, McElvain, and Lac 2011; McCrea and Horn 2006; Brodal 1954), and the presence of connections from the cerebellum to the integrator both directly and via the vestibular nuclei (McCrea and Horn 2006; Straka et al. 2006). Unlike many other cerebellar-mediated learning tasks, in the current proposal there is no role for olivary inputs and visual signals. Instead, the proposed teaching signal is created by appropriate filtering of burst and integrator signals through cerebellar dynamics such that the signal output of the cerebellum eventually equals the real eye position.

In the current instantiation, the form of the adapted teaching signals and the values of the key parameters were built in (i.e. the scaling of the saccade command driving the internal cerebellar activity, 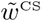, and the scaling that transforms the maximal VPNI firing rates into a maximal eye angle or muscle force, *ϑ*_max_*/r*_max_). In future variants, we anticipate that the cerebellar model could itself be calibrated through proprioceptive signals that are present beginning early in development (Graves, Trotter, and Frégnac 1987) and provide information on oculomotor muscle tension (A. Fuchs and Kornhuber 1969; Donaldson 2000), even though these proprioceptive signals are not involved in the moment-to-moment stabilization of eye position (Guthrie, Porter, and Sparks 1983; Lewis et al. 2001). Furthermore, future variants will also need to allow for potential differences in the role of cerebellar feedback during development and in the adult, at which state cerebellar output signals are largely velocity dominated (Pastor, Cruz, et al. 1997; Lisberger and A. F. Fuchs 1978).

### Cerebellum as an inverse model

In line with recent cerebellar theories (Kawato 1999; Wolpert, Miall, and Mitsuo 1998) we suggest that the cerebellum is involved in an inverse model of the plant, with the plant acting forward by converting the abducens output to the muscle force that generates the true saccade and holds the eye position. Inverting the plant implies producing from the desired saccade the appropriate abducens activities via a distributed VPNI code. The cerebellum delivers the teaching and correcting signals for the VPNI, the abducens and for itself to produce the correct input to the plant. Sequentially executing the inverse model followed by the forward action of the plant generates from the desired saccade the plant command and out of this the true saccade while suppressing inappropriate eye movements. The cerebellum together with the brainstem nuclei, including the deep cerebellar output nuclei, form an exact inverse model of the plant if the true saccade is identical the desired saccade expressed in terms of the saccade command.

The cerebellum, having phylo- and onto-genetic knowledge of the plant and an appropriate internal scaling of the saccade command, can teach the abducens neurons to produce the correct motor inputs, and the VPNI neurons to produce a robust integrator code that drives the AN neurons. Due to the nonlinearity of the plant, these teaching signals must depend on the eye position that, intriguingly, is estimated by the cerebellum based on the VPNI activities. To judge whether the CE correctly estimates the eye position, and whether the VPNI correctly integrates the saccade commands, the CE compares the own VPNI- and Saccade-driven activity (*r*^*C*^) with its low-pass filtered version 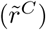. Deviations from this cerebellum-intrinsic comparison adapt the incoming synapses (from VPNI and Sacc) to improve the cerebellar estimate. A better cerebellar estimate in turn improves the teaching of the brainstem integrator circuits and the plant command (i.e. the VPNI-to-VPNI, the VPNI-to-AN, the Sacc-to-VPNI and Sacc-to-AN synapses, see Fig. 7A).

### Learning with robustness

Our developmental model also incorporates two robustness mechanisms that have been considered important for integrator function. A well-known challenge faced by integrator circuits and related analog memory networks is the problem of fine tuning – small variations in synaptic or cellular parameters can lead to severe changes in the integration or memory time scale. To overcome this challenge, various mechanisms for increasing robustness have been proposed including synaptic facilitation, dendritic bistability, and negative derivative feedback (Seung 1996; Koulakov, Raghavachari, et al. 2002; Goldman et al. 2003; Loewenstein and Sompolinsky 2003; Lim and Goldman 2013; Sanders et al. 2013). The current model naturally incorporates negative derivate feedback via a conductance-based somatic teaching input that represents a low-pass filtered version of an integrator cell’s own activity. Since the low-pass filtering smooths out the activity peaks, combining the currents induced by cerebellar input with those from local recurrent input provides a corrective difference signal that immediately dampens any drift. We also introduced synaptic depression that stabilizes population activity when stronger saccadic inputs recruits further excitatory interneurons that push the Integrator neurons to a higher sustained activity level. This synaptic depression can again be viewed as a dynamic negative derivative feedback. The developmental model learns to appropriately recruit individual integrator neurons such that, due to the synaptic depression, the sustained population activity stabilizes at a fixed set of activities, even if the teaching feedback has a systematic bias. We expect that our learning scheme is general enough to incorporate additional robustness mechanisms as well.

### Emergence from a minimal set of assumptions

The patterns of connectivity and the realistic tuning curves of the different neuronal classes emerged during learning guided by a minimal set of assumptions. A first assumption is that the cerebellar teaching signal to the AN compensates the muscle nonlinearities such that the effective eye position is eventually linearly encoded in the cerebellar teaching activity. The cerebellar teaching signal to the VPNI in turn should provide enough variability in the distribution of the thresholds and slopes of the integrator neurons such that the VPNI neurons themselves can learn to select the appropriate feedback signal to sustain their activity, and the AN neurons can learn the appropriate weighting to produce the required nonlinear muscle force for the linearly encoded left and right eye position. A second assumption is that the cerebellum has an internal (inverse) model of the oculomotor plant through which the signals from the integrator get filtered. This model could presumably be calibrated through proprioceptive inputs from the oculomotor muscles (Donaldson 2000).

### Bootstrapping in motor learning

Bootstrap learning may more generally underly motor learning when internal feedback is produced, as here by the cerebellum and more generally by a predictive forward model of the motor output, that helps to improve the inverse model (Kawato 1999; Wolpert, Miall, and Mitsuo 1998). Such feedback signals provide information about a given motor command and its induced motor state, and the comparison between the effective and desired state allows for improving both the inverse model and the internal feedback. We have shown how such a scheme can work for the learning of an inverse model that produces an appropriate motor code for the eye position out of a saccade command. More generally, learning an inverse model based on internal or external feedback to set a target for the upcoming motor state is a strategy that may also underly imitation learning (Giret et al. 2014). Internal feedback may also be provided by the recall of past errors from an error memory that helps to quicker adapt to motor perturbations (Herzfeld et al. 2014). Delays in the prediction of the feedback may be overcome by a prospective code of the cerebellar activities that can also be expressed in terms of dendritic predictive learning (Brea et al. 2016). The suggested bootstrap learning may further inspire the learning of position or movement self-stabilization in the field of robotics, where a greater need for biologically-inspired algorithms has been noted (Pfeifer, Lungarella, and Iida 2007).

## 4 Methods

### Learning to integrate using feedback

Here we formally construct the recurrent integrator network with activity *r*^V^ that integrates saccade commands *r*^S^. The basic idea is to consider a low-pass filtered version *r*^C^ of *r*^V^ that also receives the saccade inputs, and to adapt the various synaptic strengths such that eventually *r*^V^ = *r*^C^. Being equal to its low pass-filtered version is only possible if *r*^V^ is constant in between the pulse-like saccade inputs.

The general form of the recurrent integrator dynamics is (Fig. 2A)

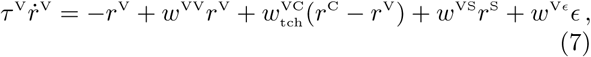

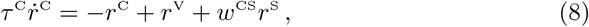

with some unbiased, quickly changing noise *E* (see Supp. Information). The time constants were set to *τ* ^V^ = 10 ms and *τ* ^C^ = 50 ms. As *r*^C^ and *r*^V^ represent firing rates, we truncated them at 0 so that they stay non-negative and we also truncated them at a maximal value (*r*_max_ = 150 Hz) to prevent running-away for initial feedback weight *w*^VV^ > 1.

The dynamics of the synaptic strengths is defined to descend the error function 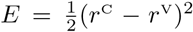 that eventually leads to *r*^V^ = *r*^C^. As according to Eqs 7 and 8 the activities depend linearly on the synaptic weights, the gradient ascent for the recurrent weight, for instance, leads to the dynamics

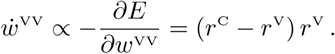

With the corresponding gradient calculation for the synaptic weights mediating the saccade inputs (and below also for the other synaptic weights) we obtain the downhill (and here even gradient) dynamics

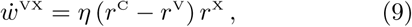

with presynaptic identity X ∈ {V, S}, a small learning rate *η* = 0.5 (and an update every Euler step, with *dt* = 1 for the one-compartmental neuron simulations). We set *w*^CS^ = *τ* ^C^ to fix the scale of the external input and also fixed 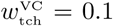. While *r*^V^ → *r*^C^ the plastic weights converge to *w*^VV^ →1 and *w*^VS^ → *τ* ^V^ respectively. This is because the steady state 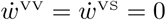 requires *r*^V^ ≡ *r*^C^, and this implies that also the dynamics of *r*^V^ and *r*^C^, i.e. Eqs 7 and 8, are identical. We conclude that *w*^VV^ = 1 and *w*^VS^ = *τ* ^V^. In this case, Eqs 7 and 8 reduce to the single integrator dynamics 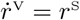 (see Supplementary Information S1.1).

### Multistability through nonlinear feedback

To improve the stability of the integrator we consider a nonlinear feedback *ψ*(*r*^V^), extending Eq. 7 to

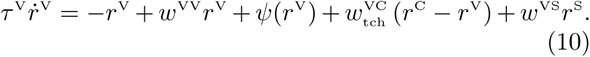

After learning one has 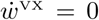 and hence *r*^C^ = *r*^V^. Correct temporal integration of *r*^S^ is obtained when (after learning) *w*^VS^ = *τ* ^V^ and when

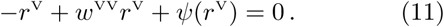

Strict equality dictates that the feedback function should be linear, *ψ*(*r*^V^) = *r*^V^(1 − *w*^VV^), but this yields a line attractor that is again sensitive to the mistuning of the recurrent feedback. Stability in turn requires the condition

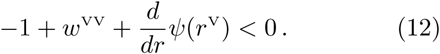

to be satisfied at least at a few rates. As a compromise between Eqs 11 and 12 we may choose *ψ*(*r*^V^) = *γ* sin(*ω r*^V^) with *γ* = .01 and *ω* = *π/*3, see Fig. 4B,C. We next show that such a nonlinearity can be learned by the same learning rule as before and further improves stability.

### Nonlinear feedback through recruitment

To learn the nonlinear feedback we consider a population of *N*_I_ = 200 interneurons (putatively excitatory or inhibitory, see also Fig. 5E) that project through short-term depressing synapses onto the principal neurons *N*_V_ = 2. The variable *r*^V^ is now expressed as the population rate 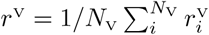, where each of the 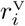 follows the dynamics in Eq. 10. The total interneuron feedback becomes

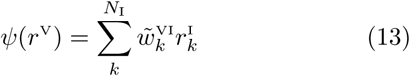

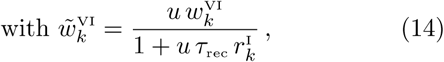

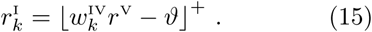

Here, 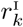 is the firing rate of the *k*’th interneuron, using ⌊*x*⌋^+^= *x* for *x* > 0 and ⌊*x*⌋^+^ = 0 else, *ϑ* = .15 is the activation threshold, and 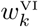 is the non-depressed synaptic weight to be learned. Synaptic depression is quantified by the recovery time constant *τ*_rec_ = 0.1 s and the ‘use’ parameter *u* = .2, giving the effective weights 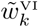 of Eq. 14 (Tsodyks and Markram 1997). The synaptic release rate becomes 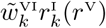 that is a saturating function of *r*^V^ (Figs 5C and 6).

A learning rule for the weights from the inter-to-principal neurons that also descends the error function *E* is obtained as the ones for the other weights (Eq. 9),

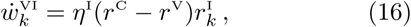

here with the learning rate *η*^I^ = 0.1*/N*_I_. Without constraining the sign of the weights some become excitatory and some inhibitory in order to generate the appropriate nonlinearity (cf. Fig. 5D). The weights from the principal to the interneurons, 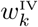, are fixed and sampled from the distribution (Fig. 5B)

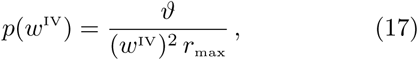

evaluated for 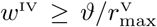. As shown in the Supp. Information S1.2, this weight distribution supports a linear recruitment of the *N*_I_ interneurons.

### Two-compartment neuron model

We next interpret 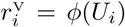 as instantaneous Poisson firing rates of 2-compartment VPNI principle neuron, with *U*_*i*_ being the somatic voltage and a threshold-linear transfer function *ϕ*(*U*) = ⌊*γU*⌋ ^+^ for some constant *γ*> 0. The somatic compartment receives a teaching current *I*_tch,*i*_ from the cerebellum. The dendritic compartment with voltage *V*_*i*_ is driven by recurrent feedback from the principal- and interneurons and by the external stimulus (Fig. 5A). As we will show, the neuronal dynamics in Eqs 8 and 10 can be obtained from the coupled dynamics of a population of VPNI neurons and the cerebellar activity,

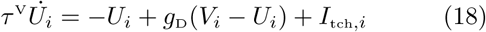

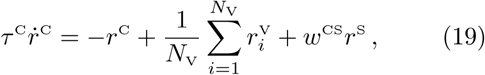

with

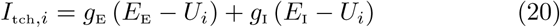

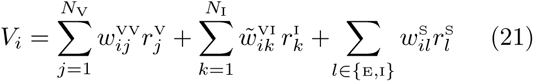

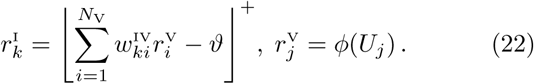

The effective weights 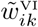 from the interneurons to the principle neurons are defined as in Eq. 14 and include synaptic depression. The interneuron transfer function is threshold-linear (Aksay, Baker, et al. 2000), with the same threshold as in Eq. 15 and the distribution of 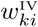 as described above. The dendritic transfer conductance and the inhibitory conductances are fixed to *g*_D_ = *g*_I_ = 20 and the excitatory and inhibitory reversal potential were set to *E*_E_ = 4⅔ and *E*_I_ = −⅓. The excitatory and inhibitory conductances *g*_E_ and *g*_I_ together define the target (‘matching’) potential *U*_M_ for the somatic potential *U*_*i*_, where

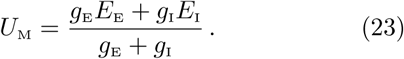

Adapting the weights on the dendritic compartment will push the somatic potential towards its target such that eventually *U*_*i*_ = *U*_M_, or in terms of spike rates, 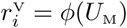. To ensure that these rates also match the cerebellar teaching rate, *r*^C^ = *ϕ*(*U*_M_), we plug (23) into this last equation and solve for *g*_E_,

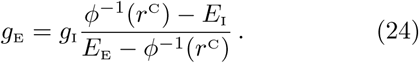

As required by the biology, *g*_E_ is a non-negative, montonically increasing function of the presynaptic firing rate *r*^C^. The analogy of the somatic teaching current in Eq. 18 to the teaching signal 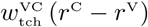 in Eq. 10 becomes apparent by rewriting (20) as

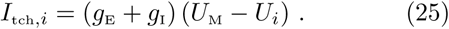

To map the dynamics for 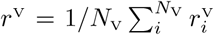 and *r*^C^ obtained from the 2-compartment model (Eqs 18 – 22) to the dynamics of the simplified model (Eqs 8 and 10) we first consider the case of one single VPNI principal neuron (*N*_V_ = 1) for which 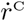 is identical in both models (Eqs 8 and 19). Since we are interested in the stabilization of the dynamics between saccades we also assume *r*^S^ = 0. The first steady state condition, 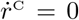, is then reached if and only if *r*^C^ = *r*^V^. Since *r*^V^ = *ϕ*(*U*) and *r*^C^ = *ϕ*(*U*_M_) we conclude that *U* = *U*_M_, and hence *I*_tch_ = 0 (Eq. 25). With this, the second steady state condition, 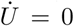 (Eq. 18, again for a single neuron, hence droping the index *i*), yields (also using Eqs 21 and 13)

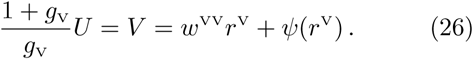

The above Eq. 26 is the condition for the steady state in the simplified model (Eq. 11) if we identify 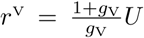. To do this we note that by construction of the VPNI and interneuron feedback the term *w*^VV^*r*^V^ + *ψ*(*r*^V^) is close to *r*^V^ and hence always positive (see Fig. 5F). We can therefore safely truncate the left-hand-side in Eq. 26 at 0 by applying ⌊. ⌋^+^. Hence, the transform (*U, U*_M_) → (*r*^V^, *r*^C^) defined by

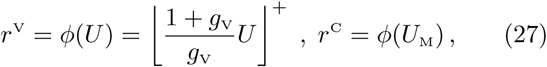

maps the steady states of the dynamics 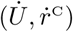 from Eqs 18 and 19 to the steady states of the dynamics 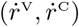 from Eqs 10 and 8, and vice verse (setting *γ* = (1 + *g*_V_)*/g*_V_). By mapping the steady states we have shown that the dynamics of the 2-compartment model is topologically equivalent to the one of the simplified model. For *N*_V_ > 1 the topological equivalence is shown by the same fixed point mapping (Eq. 27), but applied to the averaged population dynamics for the simplified and the 2-compartment model, respectively.

### Dendritic predictive plasticity

Learning adjusts the synaptic weights on the dendritic compartments such that somatic potential *U*_*i*_, driven by the dendritic potential *V*_*i*_, converges to the target potential, *U*_*i*_ → *U*_M_. The attenuated dendritic voltage that drives the neuron in the absence of teaching input is 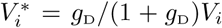. The downhill synaptic learning rules for the 3 types of synapses projecting to the principal neurons, Eq. 9, become

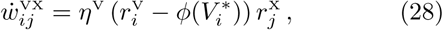

where X ∈ {V,I,S} is the identity of the presynaptic neuron. We chose *η*^V^ = 0.1*/N*_I_. Crucially, for *U*_*i*_ = *U*_M_, the somatic current vanishes (Eq. 25), and because the somatic potential becomes the attenuated dendritic potential *U*_*i*_ = *V* ^*^, the plasticity signal also vanishes (Urbanczik and Senn 2014) 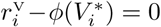. In the spiking version (Fig. 6) the presynaptic rates 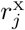 were replaced by a train of postsynaptic potentials elicited by presynaptic spike trains generated with the corresponding Poisson rates. Similarly, the postsynaptic rates 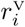 were replaced by a spike train generated from that Poisson rate. For details to Figs 5 and 6 see Supp. Info. S1.5.

### Cerebellum learning an internal model

To model the complete VPNI we consider 2*N*_V_ neurons, with the first *N*_V_(= 100) being the ipsi and the second *N*_V_ the contralateral neurons. We also consider an internal CE variable 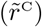 that represents the low-pass filtering of the cerebellar activity *r*^C^ plus the saccade input. Eq. 8 is then replaced by the two dynamical equations

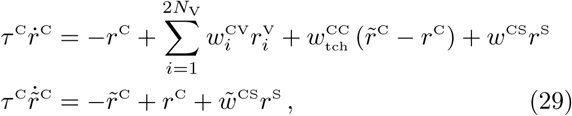

with the cerebellar self-teaching weight fixed to.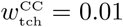.

The weights mediating the VPNI and saccade inputs, 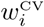 and *w*^CS^, are adapted such that *r*^C^ closes up to 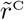,

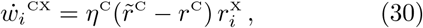

with X ∈ {V, S} and learning rate *η*^C^ = 0.1*/N*_V_. In the presence of VPNI input, CE-learning pushes *r*^C^ towards 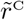. After learning, the cerebellar activity is equal to its low-pass filtered version, 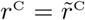, and according to Eq. 29 the CE, in the presence of the VPNI input, also integrates the saccadic inputs, 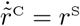 (provided that 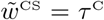 and the coefficient in front of *r*^C^ is 1). Hence, after learning the CE extracts the desired eye position 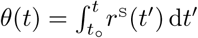 from the VPNI activity and the saccadic inputs, *r*^C^ = *θ*. Note that the CE is not an integrator when decoupled from the VPNI.

### Adaptive CE teaches VPNI

To obtain the fully self-organizing integrator circuitry we consider the adaptive cerebellum that teaches AN and VPNI neurons. The VPNI neurons are recurrently connected within and between the ipsi- and contralateral pool and receive the CE teaching signal and the saccade commands,

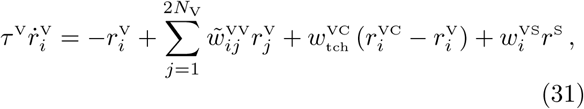

where 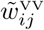 represents the depressed weight analogously defined as 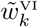 in Eq. 14 but for the presynaptic (principle) neuron with rate 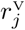. The threshold-linear transfer function is imposed by clipping the rates at 0, ensuring that *r*^V^ ≥ 0 (see Supp. Fig. S4B, applied also to the CE and AN neurons). Initially, all these weights are randomly chosen from a Gaussian around 0 with variance of 0.01 (except for a non-adaptive teaching weight that is fixed to 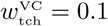, Eq. 31), as are all the other plastic weights governed by Eq. 30 (and 39 below).

The teaching signal from the cerebellum to both pools is each composed of an ‘ipsilateral’ contribution (threshold-linearly increasing with *r*^C^, *i* = 1…*N*_V_) and ‘contralateral’ cerebellar contribution (threshold-linearly decreasing in *r*^C^, *i* = *N*_V_ + 1, …, 2*N*_V_),

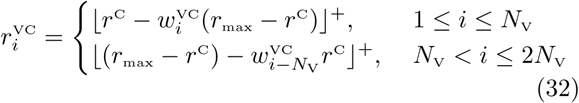

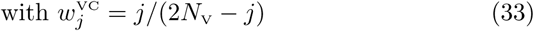

and *r*_max_ = 150 Hz. Note that for the ipsilateral VPNI pool (*i* = 1…) the excitatory teaching signal *r*^C^ can be interpreted as an ‘ipsilateral’ cerebellar contribution, and the inhibitory signal proportional to (*r*_max_− *r*^C^) as a contralateral contribution, with the symmetrical interpretation for the contralateral VPNI pool. The synaptic weights within and onto the VPNI adapt according to the standard predictive plasticity,

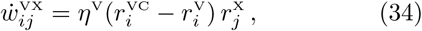

with X∈ {V, S}. For more simulation details to Fig. 7 see Supp. Info. S1.5.

The teaching signal in Eq. 32 is constructed such that during learning (where *r*^C^ → *θ*) the VPNI transfer functions converge towards 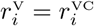 with

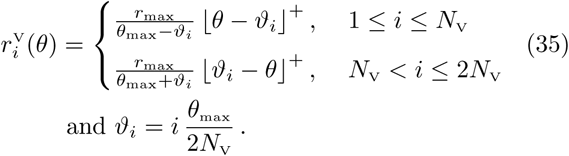

Here, *θ*_max_ = *r*_max_ (= 150 Hz). These functions roughly match the experimental ones reported in (Aksay, Baker, et al. 2000), see also Figs 7F and S2. In these figures the eye position *θ* is mapped to an eye angle via *θ* → 15° (2*θ/θ*_max_ − 1).

### Adaptive CE teaches AN

We next introduce a (ipsilateral) population of *N*_A_(= 20) abducens neurons that receive (ipsilateral) VPNI inputs, teaching input from the CE and saccade inputs,

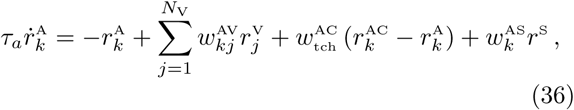

with *k* = 1 … *N*_A_. To obtain a positive threshold-slope correlation, the teaching input to the k’th AN neuron is composed of a common excitation (proportional to the ‘ipsilateral’ CE activity *r*^C^) and an individualized inhibition (proportional to the ‘contralateral’ CE activity *r*_max_− *r*^C^),

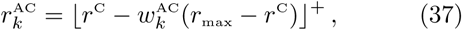

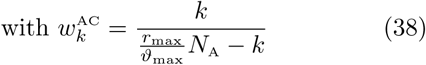

and *ϑ*_max_ = 135 Hz. The weights in Eq. 38 are chosen such that the recruitment thresholds of the teacher neurons are uniformly distributed. The synapses projecting from VPNI and Sacc to AN are adapted by

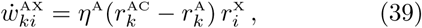

with X ∈ {V, S}, learning rate *η*^A^ = 0.1*/N*_AX_ and teacher signal 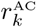 given by Eq. 37. Learning of these synapses implies that 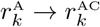 (Figs 7E and S3).

The AN neurons are assumed to elicit a total muscle force 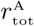 of the lateral eye muscle that is a weighted sum of the individual AN firing rates, 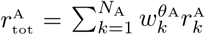, with weights 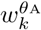 that increase with *k* and thus with the recruitment threshold as described by the size principle (McFarland and Albert F. Fuchs 1992), see Supplementary Information S1.3. The produced eye position as a function of the total muscle force is set to

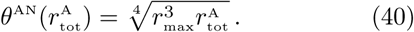

This function was derived such that, after learning, the produced eye position fits its cerebellar representation, *θ*^AN^ ≈ *r*^C^. Because the latter also matches the desired eye position, *r*^C^ ≈ *θ*, we eventually get *θ*^AN^ ≈ *θ*.

### Animal rearing

All experiments were approved by Weill Cornell Medical College Institutional Animal Care and Use Committee. Wild-type larval zebrafish (Danio rerio) embryos from the same clutch were separated into two groups at 1–4 hours post fertilization. One group (n=13) was reared under a 12 hour light/12 hour dark cycle while the other (n=18) was reared in an opaque encasing under a 24 hour dark cycle. All larvae were kept at 28°C in a thermostat-controlled incubator.

### Preparation and eye movement recordings

5 day post fertilization larvae were anesthetized in a 200 *ng/µ* L solution of MS-222 and embedded in 1.2% low melting-point agarose (Sigma-Aldrich, A0701-100G). Agarose was removed from around the eyes to allow for undamped eye movements. Animal mounting and agarose removal were carried out in less than 5 minutes to minimize exposure to light in dark reared animals. Animals were given two hours to wake up from anesthesia (dark reared animals were kept in the dark during this time) and then placed in the recording chamber. Animals were illuminated using infrared LEDs (850 nm, ThorLabs) not visible to the zebrafish in order to record eye movements made in the dark. Animals were given five minutes to acclimate to the recording chamber. Since adult zebrafish sometimes show reduced spontaneous behavior after being in the dark for too long (Mensh et al. 2004), we exposed all animals to one minute of visible light prior to recording. Spontaneous eye movements were recorded in the dark for five minutes at 25 Hz using a sub-stage infrared CMOS camera (Allied Vision Technologies, Guppy FireWire camera). Eye position was extracted from recordings by fitting an ellipse (using the Matlab command regionprops) to contrast-thresholded images as described in (Beck et al. 2004).

### Data Analysis

We only analyzed data from animals that showed healthy eye movement development as measured by saccade frequency and amplitude. We did not include data from animals that made fewer than six saccades over a five minute interval or from animals that only made saccades in one direction (each animal needed to make at least two saccades in both directions over a five minute interval). Each inter-saccade interval was required to be less than 100 seconds for the dataset to be included. In addition, we did not include data from animals that showed a restricted range of movement (amplitudes less than 10 degrees). Ocular drift during fixations was measured by binning eye position into 500 ms segments and calculating the best fit slope describing eye position versus time within each bin. PV plots were generated by plotting best fit slopes versus average eye position within bins (Major et al. 2004). We excluded the first second following each saccade from our analyses to avoid transient eye movement dynamics such as post-saccade slide that do not solely reflect neural integrator output. We also excluded fixations following swim-like movements and eye movement twitches.

## Acknowledgment

This work is dedicated to the late Robert Urbanczik with whom we (EN and WS) shared many exciting scientific discussions and whom we keep in memory for his unique and illuminating perspective on science and life in general.

## S1 Supplemental Information

### S1.1 Solving the 2-dimensional integrator system

In the absence of external input, Eqs 7 and 8 form a linear homogeneous system that has the solution

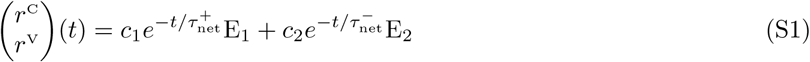

with constants *c*_1_ and *c*_2_ depending on the initial values, and network time constants 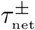 specified by

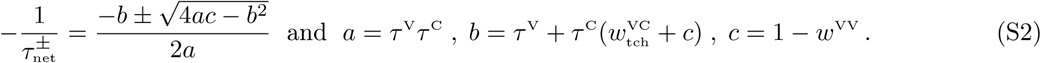

When the recurrent weight converges to unit, *w*^VV^ → 1, the long time constant tends to infinity 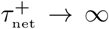 and the corresponding eigenvector E_2_ converges to the unit vector. In this case, assuming initial conditions *r*^C^ = *r*^V^ = *c*_2_ (implying *c*_1_ = 0), the inhomogeneuous system becomes a perfect integrator,

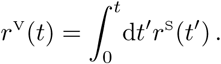

The slow time const 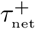 is shown in Figs 2E and 3B.

### S1.2 Heavy-tail weight distribution providing linear recruitment

Since the synaptic release rate is saturating with increasing presynaptic rate *r*^V^ due to synaptic depression (cf. Fig. 5C), the relevant quantity is the number of interneurons *n*(*r*^V^) that crossed the threshold *ϑ* at a certain rate *r*^V^ of the population rate of the principal neurons. A neuron becomes active as soon as its total synaptic drive crosses threshold, 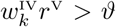. If *p*(*w*) and *P*(*w*) denote the density and cumulative density of the V → I weights 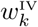, respectively, we get

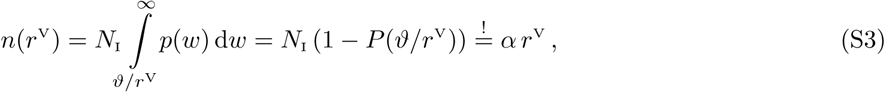

where the lower limit of the integral, *ϑ/r*^V^, represents the minimal total weight *w* needed to activate a first interneuron given the presynaptic rate *r*^V^.

Assuming that at the maximal presynaptic firing rate 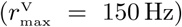 all interneurons are recruited, 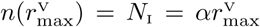, we infer that 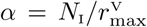. From Eq. S3 we then get the self-consistency relation 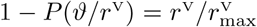 that, substituting *ϑ/r*^V^ = *w*, leads to 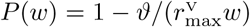. Hence, the density that leads to a linear recruitment (Fig. 5B) is

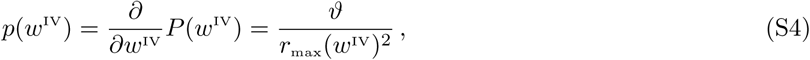

that is evaluated for values 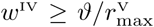 according to Eq. S3. For each interneuron *k* we sampled the total excitatory weight 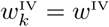 according to the density in Eq. 17 and randomly split up this total weight into *N*_V_ individual weights 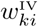 constrained to 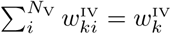. Remarkably, the statistics of the individual weights 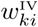 across *i* and *k* nicely fits a log-normal distribution (Fig. 5B), in agreement with experimental studies (Koulakov, Hromádka, and Zador 2009). Simulations with a population of such interneurons shows enhanced stability after learning, even in absence of the stabilizing delayed feedback (Fig. 4F).

### S1.3 Eye position produced by AN that matches CE activity

Setting 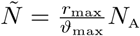 and plugging the weights from Eq. 38 into Eq. 37 we obtain

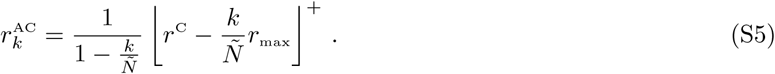

After successful learning, the firing rate of the AN neuron driven by the VPNI input becomes equal to the teaching signal from the CE to that same AN neuron, 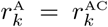. Choosing the weight that specifies the contribution of the *k*’th AN neuron to the total muscle force as 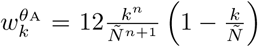 for *k* = 1 … *N*_A_ and *n* ≥ 0 we then calculate,

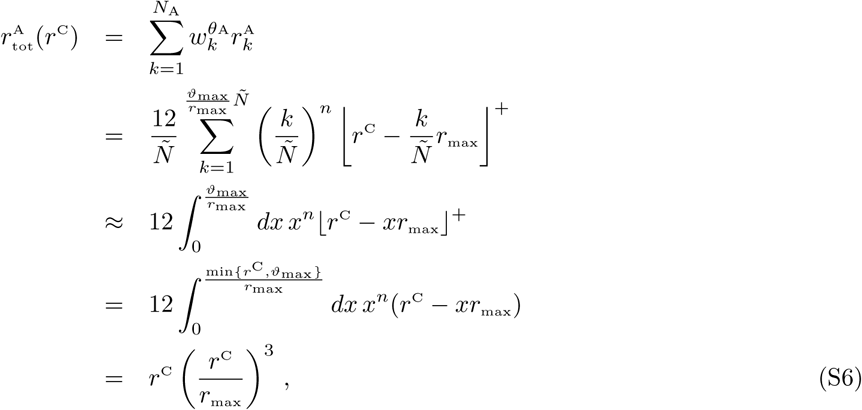

where the approximation is valid for large *N*_A_ and where in the last step we set *n* = 2 and assumed *r*^C^ ≤ *ϑ*_max_. Solving this relation for the cerebellar activity yields 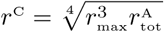. Hence, defining the eye position produced by the total muscle force 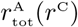 as

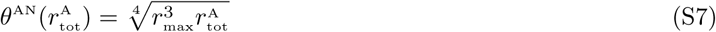

gives an eye position that matches the cerebellar activity, *θ*^AN^ = *r*^C^, for *r*^C^ ≤ *ϑ*_max_ (see Fig. S3D).

Note that the readout weight 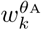 defined above are maximal for 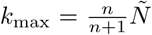. The choice of *n* = 2 with the given values of *r*_max_ (= 150 Hz) and *ϑ*_max_ (= 135 Hz) is motivated by fact that the maximum is reached for 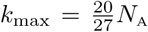, and hence the weights are growing with *k* = 1 … *N*_A_ up to the highest indices. Put in other words, with the final distribution of AN firing properties (see Fig. S3D and (Pastor, Torres, et al. 1991)) and a force-to-position function 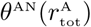 that shows at least a 4’th square root saturation, the size principle must emerge to compensate for the overall nonlinearities from the cerebellar input to the eye position (McFarland and Albert F. Fuchs 1992). In the cat, the muscle force as a function of the eye eccentricity (the analog to Eq. S6) was in fact measured to be exponential (Carrizosa et al. 2011), favoring the size principle (Carrascal et al. 2011).

### S1.4 Simulation details for Figures 2, 3 and 4

**Figure 2** We modified the coupled differential equations by adding a noise term ϵ to the integrator neuron *r*^V^

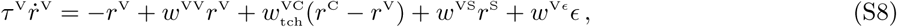

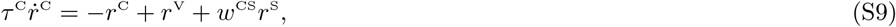

where the noise variable *ϵ* is sampled from an Ornstein-Uhlenbeck process with 1*/θ* = 5 ms, *µ* = 0, *σ* = 1 and with weight *w*^V*ϵ*^ = 0.005 was chosen such that the noise was approximately 5% of the signal amplitude. The plastic weights (*w*^VV^, *w*^VS^) were initially drawn from a normal distribution 𝒩(0, 0.01) in order to remove initial setup bias (*w*^CS^ = *τ*^C^). Time courses were averaged across 20 trials (standard deviation as light green shaded area). In all the Figures the saccades *r*^S^ were modeled as jumps at random times with uniformly sampled amplitudes from a discrete set of values (typically 10), with exponential decay with time constant 10 ms. The amplitudes were chosen such that the eye position *θ*^AN^ (Eq. S7) roughly lies between 0 and 150 (with cutoff at 0). Other parameter values: *τ* ^V^ = 10 ms, *τ* ^C^ = *w*^CS^ = 50 ms.

**Figure 3** Simulation of the same equations as in Figure 2 was used with the same parameters for the noise and the same value for the other parameters, unless indicated differently. The effective time constant of the system was derived analytically (Sec. S1.1) by looking at the eigenvalues of the coupled dynamical equations (see Eq. S2).

**Figure 4** We simulated the dynamical equations 5 & 6 with an additional noise term weighted as in Equation S8. All plastic weights were initially drawn from a normal distribution 𝒩(0, 0.01) in order to remove initial setup bias. A simulation timestep of 1*ms* was sufficiently small for the given dynamics.

### S1.5 Simulation details for Figures 5, 6 and 7

**Figure 5** After learning, in the absence of teaching and saccadic input, the steady state of the somatic compartment (Eq. 18, −*U*_*i*_ + *g*_D_(*V*_*i*_ − *U*_*i*_) = 0) arises due to the balance of excitatory and inhibitory in- and outflux,

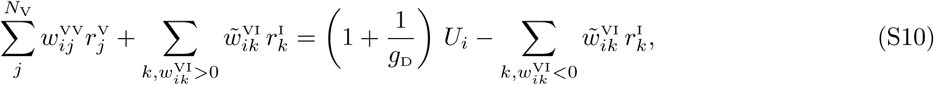

where the left-hand-side of the equation contains the recurrent excitation and the excitatory portion of the interneuron input (red trace in Fig. 5D). The right-hand-side represents the inhibitory contribution (with negative weights from interneurons to principal neurons), together with the effective leak (blue trace in Fig. 5D) of the equation. This equality needs to hold to keep the temporal integral of the external inputs between saccade commands.

**Figure 6** For the spiking implementation we mimicked a 10 times larger population of principal neurons by stretching the instantaneous (Poisson) rates 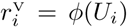 with a factor of 10 and sampled from these higher rates (while reducing the outgoing weights by the same factor). The synaptic releases train of synapse *k* is obtained from the filtered presynaptic spike train with release probability 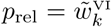, such that eventually the synaptic releases become a (inhomogenuous) Poisson train with rate 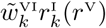, see Eqs 13–15. The simulation time step was dt= 0.1 ms. The spike-triggered postsynaptic potentials summing up the to dendritic voltage 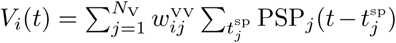, Eq. 20, have the form 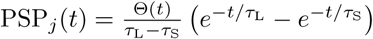 with *τ*_L_ = 10 ms, *τ*_S_ = 3 ms and Θ the unit step function. In the plasticity rule (Eq. 28) the presynaptic rates 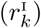 and the postsynaptic rates (*r*^V^) were replaced by spikes sampled from the corresponding instantaneous Poisson rates, while the dendritic estimate, 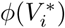, of the postsynaptic spike rate, 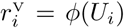, remained analogue, see also (Urbanczik and Senn 2014).

**Figure 7** The whole network of CE, AN and VPNI was simulated by running the neuronal (Eqs 29, 36, 31) and synaptic (Eqs 30, 39, 34) dynamics, driven by random saccade inputs *r*^S^ (see below). In simulations of the full model the inputs were truncated when ∫ d*t r*^S^(*t*) would have escaped the interval [0.1, 0.9] *θ*_max_. The same interval was also imposed for the CE-internal teaching signal 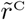.

Two different teaching protocols were applied. Continuous eye movement commands (e.g. sine-wave) were used to pre-tune the network to a line-attractor dynamics. To obtain a discrete set of stable fixed points, saccade pulses *r*^S^(*t*) were randomly applied each 200 ms or 400 ms with exponential decay (*τ* = 10 ms) and such that the integrated value (the desired eye position) 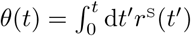 jumps to a randomly selected level out of {0, *θ*_max_*/n*, …, (*n* − 1)*θ*_max_*/n*} for a fixed *n* = 5, 10, 15 or 20, see Supplementary Fig. S5).

### S1.6 Further analysis figures

**Figure S1:**
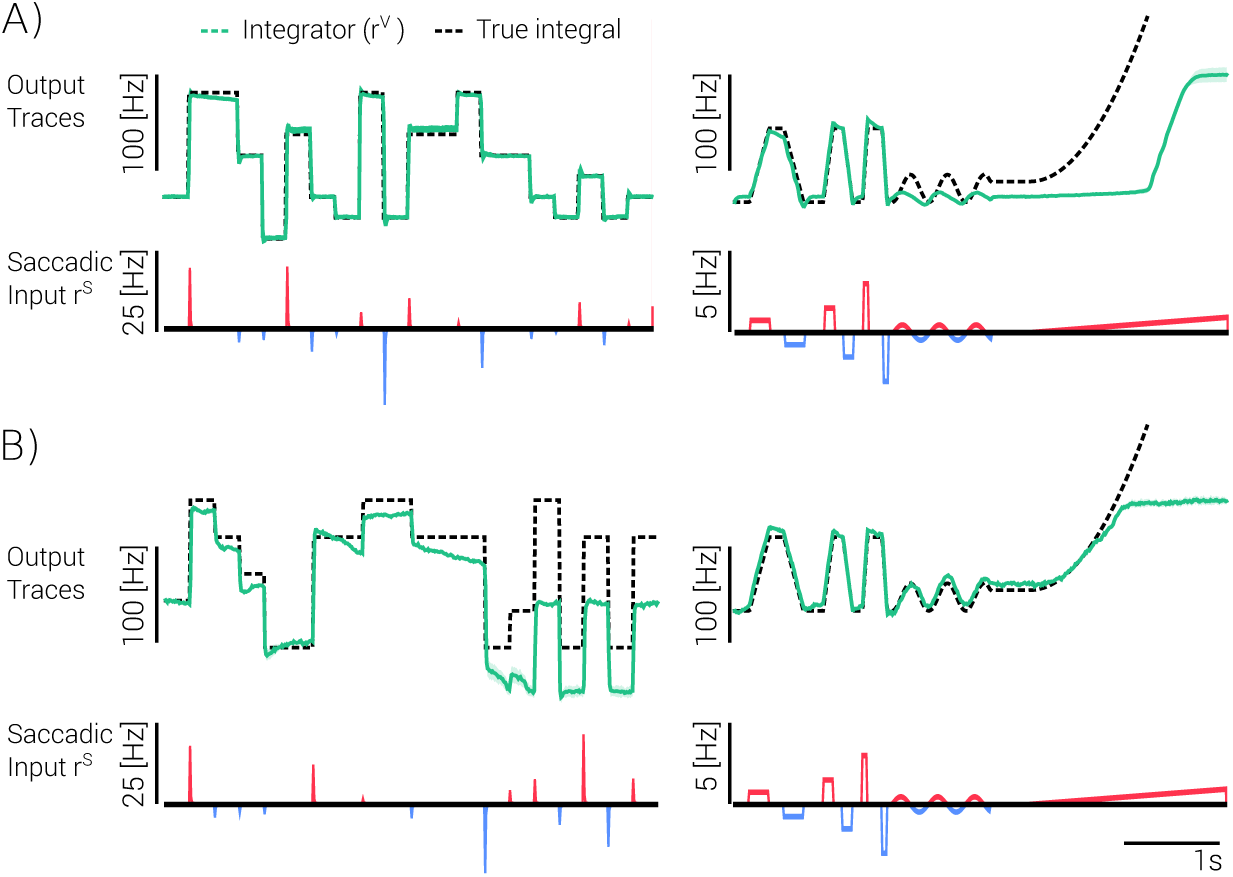
Tradeoff between capacity and robustness. The performance of the integrator depends on the stimulation protocol during learning. **A)** Performance after learning when the input r^S^(t) was chosen to be continuous followed by saccade pulses with 8 integration values (cf. Fig. S5). Left: stable response to saccade inputs. Right: response to mixed saccade and continuous input. The high stability of the fixed points learned from the saccade stimulation prevents the continuous integration of ongoing signals. **B)** Performance when the saccade inputs after the continuous stimulation was omitted. The integration of discrete pulses is not as stable as before (left), but continuous inputs with only short pauses are integrated more reliably (right).

**Figure S2:**
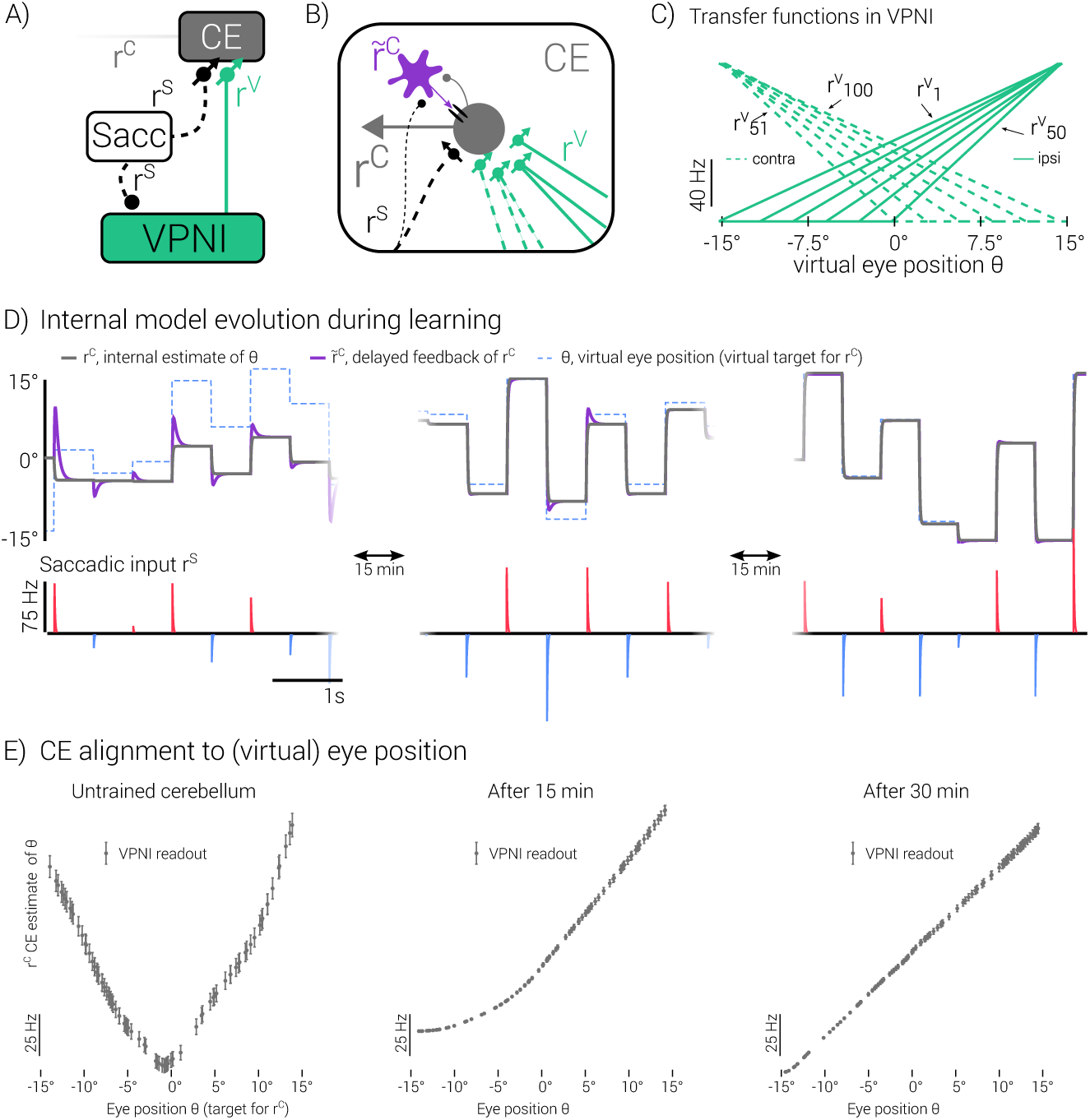
The cerebellum aligns VPNI input to its internal model of the eye. **A)** The CE and the VPNI receive common saccade commands as before, but the VPNI-to-CE connections are now also plastic. **B)** The cerebellar signal *r*^C^ adjusts itself to a fixed internal representation 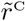 (purple) of the eye position (*θ*) through bootstrap learning (Eqs 29,30). **C)** Typical firing rates 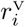 from VPNI neurons as a function of *θ* (transformed to degrees, see Eq. 35 and thereafter). These transfer functions are explicitly set here to have the shape as in the adult zebrafish, but are jointly learned in Fig. 7 via cerebellar teaching signal that recurrently builds up. **D)** During learning, the CE activity *r*^C^ (black) converges to the internal target 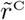 (purple) that itself converges to the integral of the saccade inputs *θ* (dotted blue line). **E)** The CE activity *r*^C^ aligns to the eye position *θ*.

**Figure S3:**
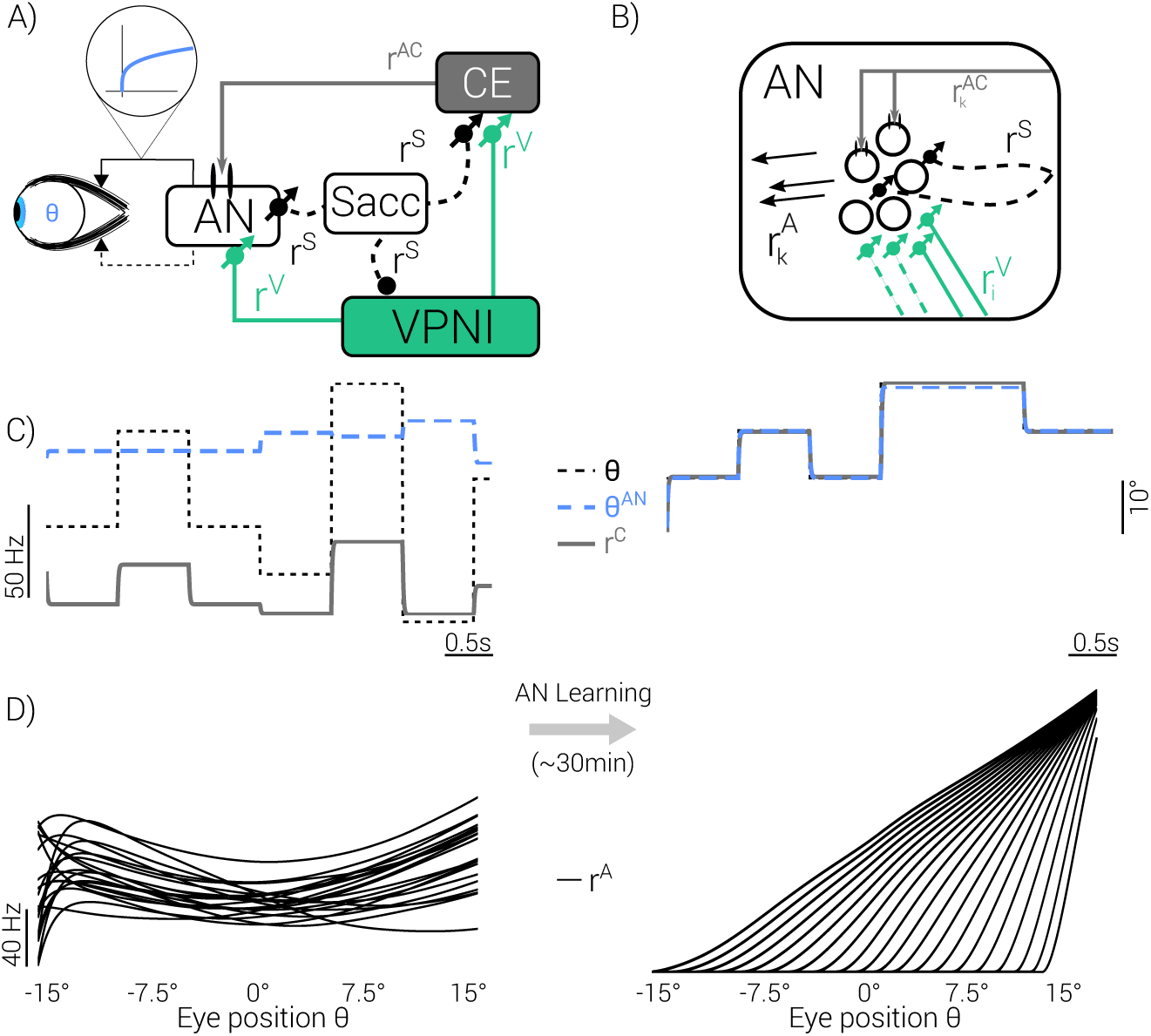
The cerebellum teaches motor neurons to produce correct eye positions from VPNI input. **A)** The cerebellar teaching signal *r*^C^ (that is matched via 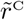 to the desired eye position *θ*, see Fig. S2) generates appropriate targets for the AN neurons. Inset: force-to-position transfer function (Eq. 40). **B)** AN neuron *k* receives teaching signal 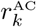 (grey) from the CE and learns to reproduce that signal out of the VPNI activity 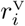 (green). **C)** The AN-produced eye position (*θ*^AN^, blue dashed), the CE-internal estimate (*r*^C^, grey) and the saccade integral (*θ*, black dots) initially differ, but match after learning (simulation of Eqs 36–40). **D)** The position-to-rate transfer functions for the AN neurons develop through learning to the ones experimentally observed (cf. (Pastor, Torres, et al. 1991)).

**Figure S4:**
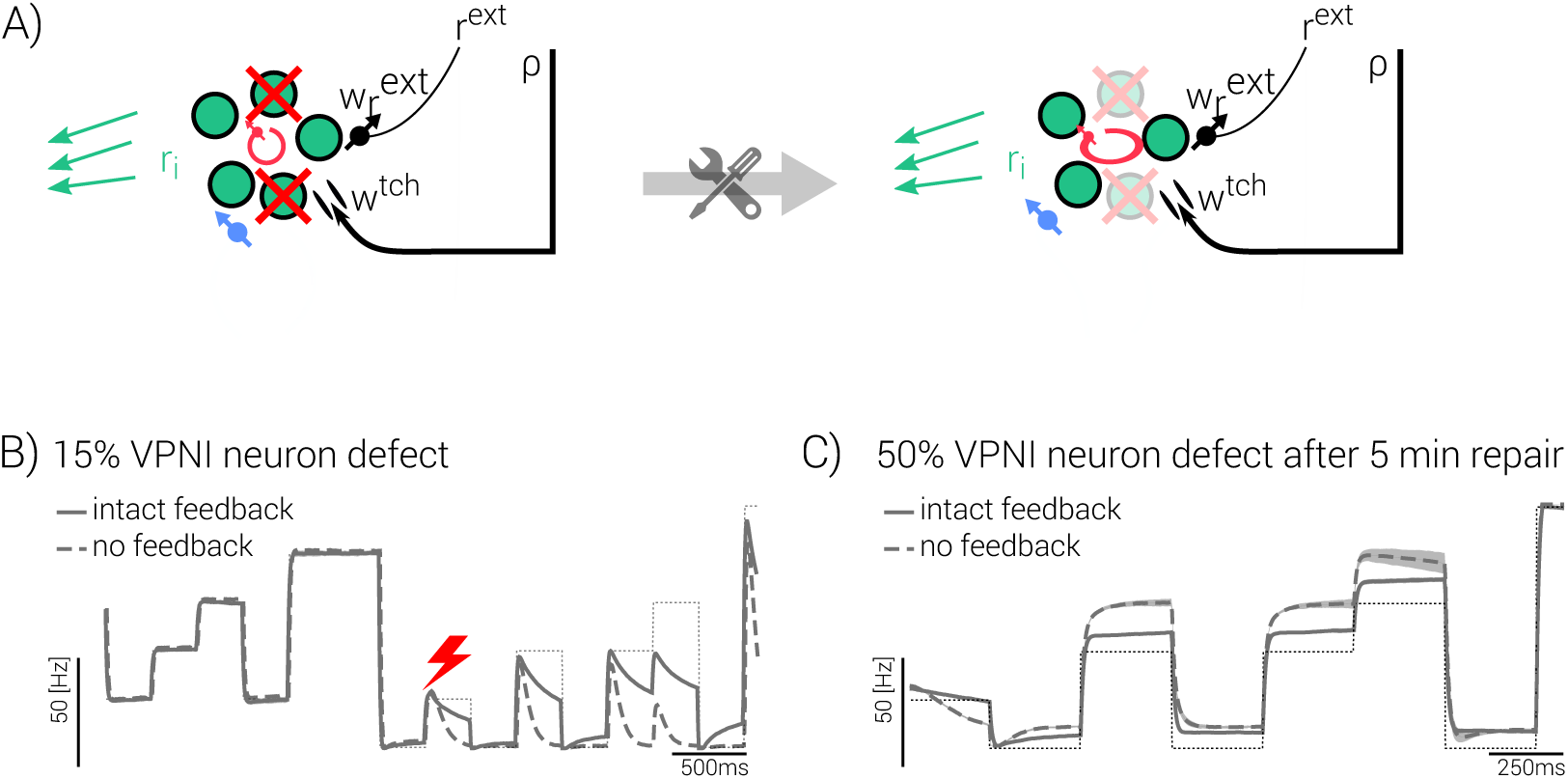
Repair after neuronal damage in VPNI. **A)** Neuronal damage within the VPNI evokes an error signal that repairs the damage based on a smaller number of neurons. **B)** With 15% fall out of the N = 100 VPNI neurons (red flash) the integration property is transiently lost and the teaching feedback immediately sets in as a correction signal (solid versus dashed). **C)** Even with 50% damage, the VPNI regains full performance after only 5 minutes. This reparation time is considerably faster than the learning from a random weight initialization that lasted more than 1 hour.

**Figure S5:**
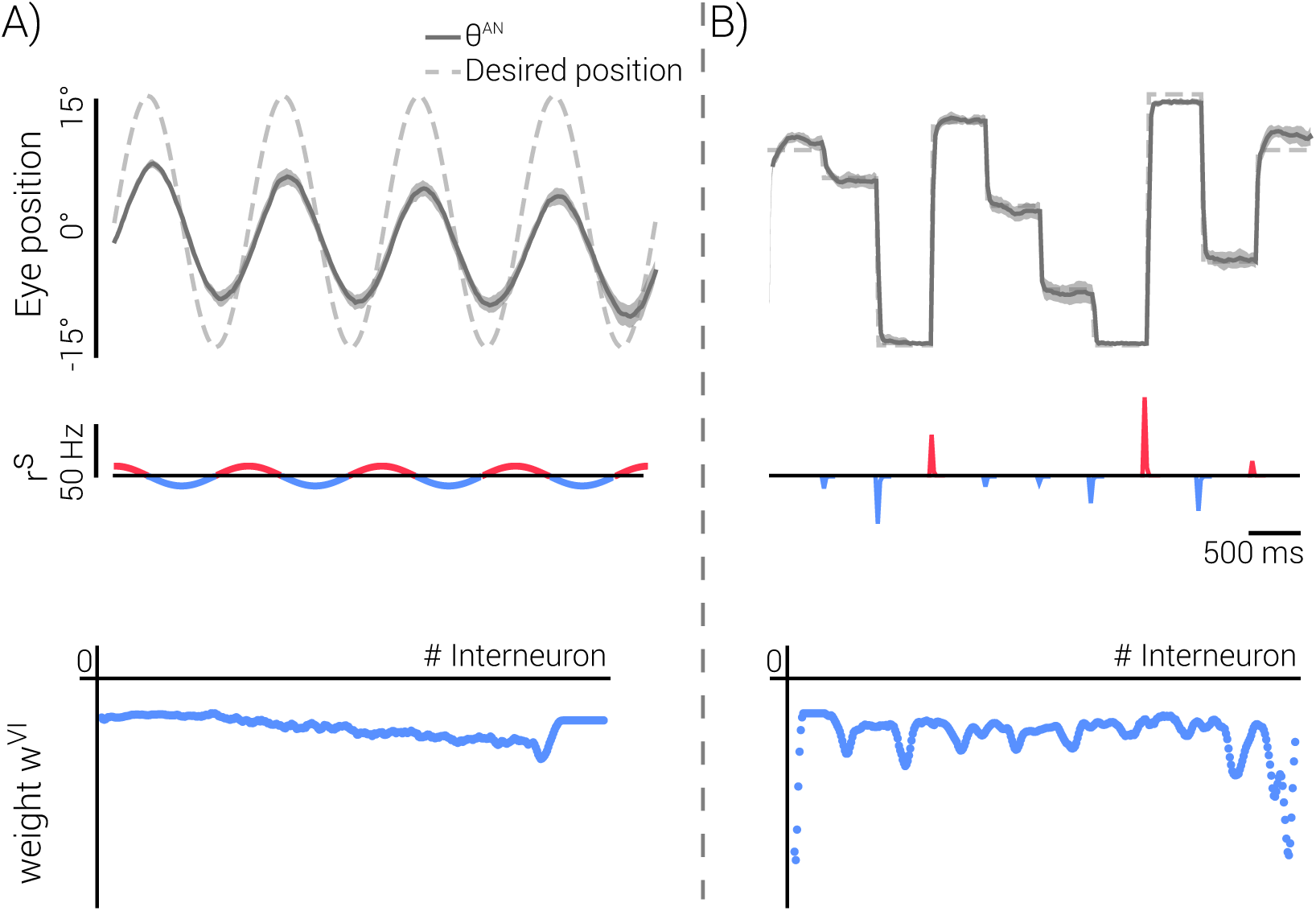
Stimulation protocol for integrator learning. Our stimulation protocol consisted of a period with continuous inputs r^S^(t) (30 min, **A**) followed by an equally long period with discrete saccade inputs (**B**, cf. Methods). This protocol mimics the natural eye movements of zebrafish larvae during the first postnatal days (Easter and Nicola 1997). Bottom row: The corresponding readout weights 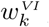 from inter-to-principle neurons in the VPNI reflects the line-attractor dynamics for the continuous input scenario (A), while it reflects the stable integration property for the saccade input scenario (B).

**Figure S6:**
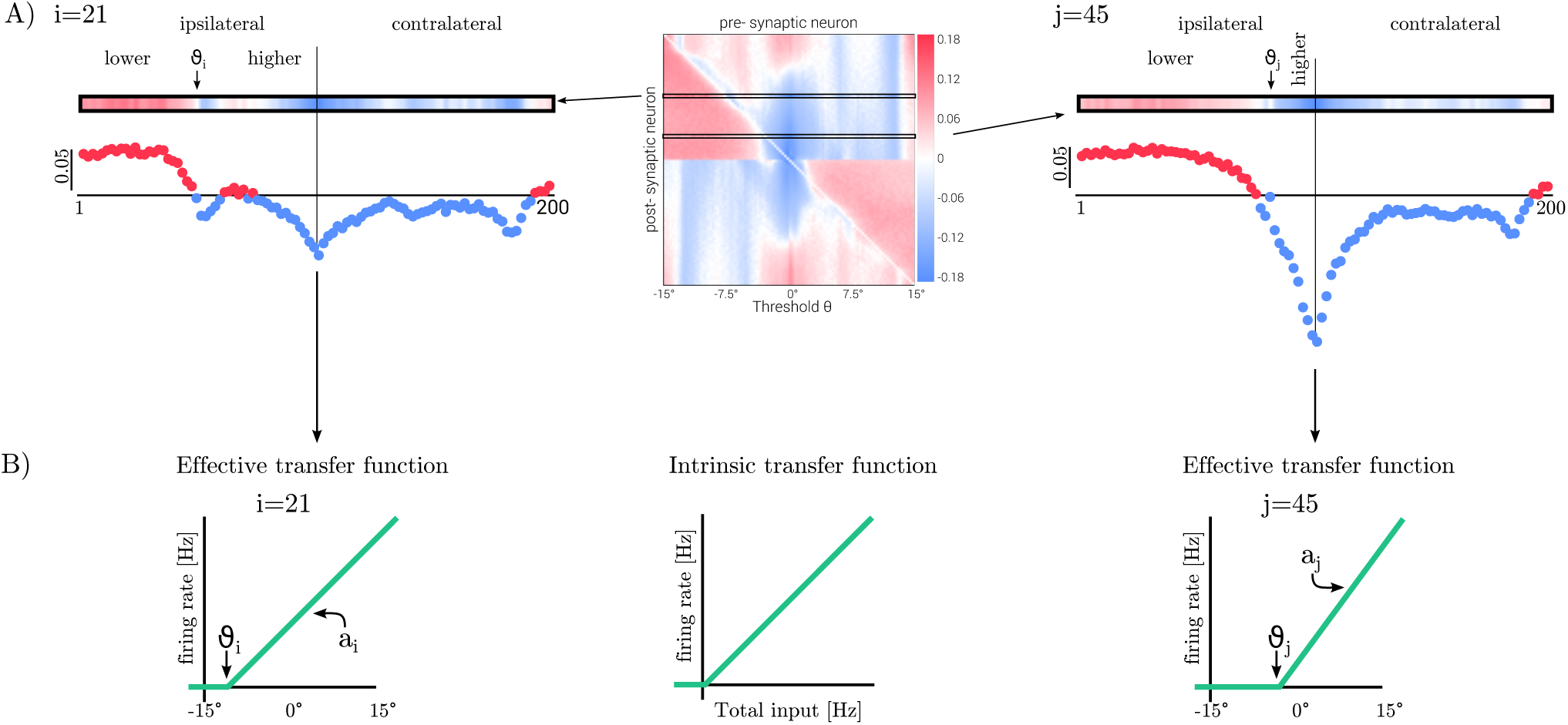
Recurrent synaptic weight structure within VPNI after learning. **A)** Recurrent weights from a presynaptic neuron (x-axis) to a postsynaptic neuron (y-axis), labeled by the eye position (θ) at which that neuron starts firing (and thus has recruitment threshold ϑ = θ). Example weight vector for postsynaptic neuron i = 21 (left) and j = 45 (right). Neurons on the ipsilateral side that have been activated before (‘lower’, with recruitment threshold ϑ < ϑ_i/j_) excite the postsynaptic neuron i/j. Ipsilateral neurons that are immediately recruited at increased eye positions (‘higher’, with recruitment threshold ϑ > ϑ_i/j_) inhibit neuron i/j, while ipsilateral neurons that are recruited later may become again moderately excitatory. Neurons on the contralateral side of i/j are typically inhibitory, up to a few neurons that are activated at very peripheral eye positions. (**B**) The intrinsic input-output transfer function (middle, with input expressed as weighted presynaptic rates, see Eq. 31) is threshold-linear and identical for all VPNI neurons (implemented by clipping the output rates at 0 to ensure 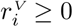). The effective transfer function as a function of the eye position is determined by the weight structure. Neuron j (right) that is recruited after neuron i (ϑ_j_ > ϑ_i_) has a steeper slope (a_j_ > a_i_) than neuron i (left), as experimentally observed (Aksay, Baker, et al. 2000). In the model this structure emerges from the cerebellar ipsi- and contra-lateral teaching signal that is excitatory and inhibitory, respectively (Eq. 32).

## References

Aksay, Emre, Robert Baker, H. Sebastian Seung, and David W. Tank (2000). “Anatomy and discharge properties of pre-motor neurons in the goldfish medulla that have eye-position signals during fixations.” In: Journal of Neurophysiology 84.2, pp. 1035–1049.

Aksay, Emre, G Gamkrelidze, H. Sebastian Seung, Robert Baker, and David W. Tank (2001). “In vivo intracellular recording and perturbation of persistent activity in a neural integrator.” In: Nature Neuroscience 4.2, pp. 184–193.

Aksay, Emre, Itsaso Olasagasti, Brett D. Mensh, Robert Baker, Mark S. Goldman, and David W. Tank (2007). “Functional dissection of circuitry in a neural integrator.” In: Nature Neuroscience 10.4, pp. 494–504.

Beck, James C., Edwin Gilland, Robert Baker, and David W. Tank (2004). “Instrumentation for measuring oculomotor performance and plasticity in larval organisms.” In: Methods in cell biology 76, pp. 385–413.

Brea, Johanni, Alexisz Tamás Gaál, Robert Urbanczik, and Walter Senn (2016). “Prospective Coding by Spiking Neurons”. In: PLoS Computational Biology 12.6, pp. 1–25.

Brodal, Alf (1954). “The cerebellar projection of the peri-hypoglossal nuclei (nucleus intercalatus, nucleus praepositus hypoglossi and nucleus of roller) in the cat”. In: Journal of Neuropathology and Experimental Neurology 13.4, pp. 515–527.

Brody, Carlos D. and Timothy D. Hanks (2016). “Neural underpinnings of the evidence accumulator”. In: Current Opinion in Neurobiology 37, pp. 149–157.

Brody, Carlos D., Ranulfo Romo, and Adam Kepecs (2003). “Basic mechanisms for graded persistent activity: Discrete attractors, continuous attractors, and dynamic representations”. In: Current Opinion in Neurobiology 13.2, pp. 204–211.

Carrascal, Livia, Jose Luis Nieto-González, Blas Torres, and Pedro Nunez-Abades (2011). “Diminution of voltage threshold plays a key role in determining recruitment of oculomotor nucleus motoneurons during postnatal development”. In: PLoS ONE 6.12, pp. 1–8.

Carrizosa, María A. Davis-López de, Camilo J. Morado-Diaz, J. M. Miller, R. R. de la Cruz, and Angel M. Pastor (2011). “Dual encoding of muscle tension and eye position by abducens motoneurons”. In: Journal of Neuroscience 31.6, pp. 2271–2279.

Denève, Sophie and Christian K Machens (2016). “Efficient codes and balanced networks.” In: Nature Neuroscience 19.3, pp. 375–82.

Donaldson, I.M.L. (2000). “The functions of the proprioceptors of the eye muscles”. In: Phil. Trans. R. Soc. Lond. B 355, pp. 1685–1754.

Easter, Stephen S and Gregory N Nicola (1997). “The Development of Eye Movements in the Zebrafish (Danio rerio)”. In: Dev Psychobiol. 31.4, pp. 267–76.

Fiorentini, A and L Maffei (1977). “Instability of the eye in the dark and proprioception”. In: Nature 270. December 15, pp. 330–331. NIHMS150003.

Fisher, Dimitry, Itsaso Olasagasti, David W. Tank, Emre Aksay, and Mark S. Goldman (2013). “A modeling framework for deriving the structural and functional architecture of a short-term memory microcircuit”. In: Neuron 79.5, pp. 987–1000. NIHMS150003.

Fuchs, A.F. and H.H. Kornhuber (1969). “Extraocular muscle afferents to the cerebellum”. In: Journal of Physiology 200, pp. 713–722.

Giret, Nicolas, Joergen Kornfeld, Surya Ganguli, and Richard H R Hahnloser (2014). “Evidence for a causal inverse model in an avian cortico-basal ganglia circuit”. In: Proceedings of the National Academy of Sciences 111.16, pp. 6063–6068.

Gold, Joshua I. and Michael Shadlen (2007). “The neural basis of decision making”. In: Annu Rev Neurosci 30, pp. 535–574. NIHMS150003.

Goldman, Mark S., Joseph H. Levine, Guy Major, David W. Tank, and H. Sebastian Seung (2003). “Robust Persistent Neural Activity in a Model Integrator with Multiple Hysteretic Dendrites per Neuron”. In: Cerebral Cortex 13.11, pp. 1185–1195.

Goldman-Rakic, Patricia S. (1995). “Cellular basis of working memory”. In: Neuron 14.3, pp. 477–485.

Graves, A L, Y Trotter, and Y Frégnac (1987). “Role of extraocular muscle proprioception in the development of depth perception in cats.” In: Journal of neurophysiology 58.4, pp. 816–831.

Guthrie, B.L., J.D. Porter, and D.L. Sparks (1983). “The functions of the proprioceptors of the eye muscles”. In: Science 16, pp. 1193–1195.

Herzfeld, David J., Pavan A. Vaswani, Mollie K. Marko, and Reza Shadmehr (2014). “A memory of errors in sensorimotor learning”. In: Science 345.6202, pp. 1349–1353.

Kawato, Mitsuo (1999). “Internal models for motor control and trajectory planning”. In: Current Opinion in Neurobiology 9.6, pp. 718–727.

Kolkman, K. E., L. E. McElvain, and S. d. Lac (2011). “Diverse Precerebellar Neurons Share Similar Intrinsic Excitability”. In: Journal of Neuroscience 31.46, pp. 16665–16674.

Koulakov, Alexei, Tomás Hromádka, and Anthony M. Zador (2009). “Correlated connectivity and the distribution of firing rates in the neocortex.” In: The Journal of Neuroscience 29.12, pp. 3685–3694. 0809.1630.

Koulakov, Alexei, Sridhar Raghavachari, Adam Kepecs, and John E. Lisman (2002). “Model for a robust neural integrator”. In: Nature Neuroscience 5.8, pp. 775–782.

Lee, M. M., Aristides B. Arrenberg, and Emre Aksay (2015). “A structural and genotypic scaffold underlying temporal Iintegration”. In: Journal of Neuroscience 35.20, pp. 7903–7920.

Lewis, R.F., D.S. Zee, M.R. Hayman, and R.J Tamargo (2001). “Oculomotor function in the rhesus monkey after deafferentationof the extraocular muscles”. In: Experimental Brain Research 141.3, pp. 349–358.

Lim, Sukbin and Mark S. Goldman (2013). “Balanced cortical microcircuitry for maintaining information in working memory.” In: Nature Neuroscience 16.9, pp. 1306–14.

Lisberger, PS. G. and A. F. Fuchs (1978). “Role of primate flocculus during rapid behavioral modification of vestibuloocular reflex. II. Mossy fiber firing patterns during horizontal head rotation and eye movement”. In: Journal of Neurophysiology 41(3), pp. 764–777.

Loewenstein, Yonatan and Haim Sompolinsky (2003). “Temporal integration by calcium dynamics in a model neuron”. In: Nature Neuroscience 6.9, pp. 961–967.

Major, Guy, Robert Baker, Emre Aksay, H. Sebastian Seung, and David W. Tank (2004). “Plasticity and tuning of the time course of analog persistent firing in a neural integrator.” In: Proceedings of the National Academy of Sciences 101.20, pp. 7745–50.

McCrea, Robert A. and Anja K. E. Horn (2006). “Morphology and Physiology of the Cerebellar Vestibulolateral Lobe Pathways Linked to Oculomotor Function in the Goldfish”. In: Progress in brain research 151, pp. 205–30.

McFarland, Jenny and Albert F. Fuchs (1992). “Discharge patterns in nucleus prepositus hypoglossi and adjacent medial vestibular nucleus during horizontal eye movement in behaving macaques.” In: Journal of Neurophysiology 68.1, pp. 319–32.

Mensh, Brett D., Emre Aksay, Daniel D. Lee, H. Sebastian Seung, and David W. Tank (2004). “Spontaneous eye movements in goldfish: Oculomotor integrator performance, plasticity, and dependence on visual feedback”. In: Vision Research 44.7, pp. 711–726.

Murakami, Ikuya and Patrick Cavanagh (1998). “A jitter after-effect reveals motion-based stabilization of vision”. In: Nature 395.6704, pp. 798–801.

Pastor, Angel M., Rosa R. De la Cruz, Robert Baker, C. I. De Zeeuw, P. Strata, and J. Voogd (1997). “Characterization of Purkinje cells in the goldfish cerebellum during eye movement and adaptive modification of the vestibulo-ocular reflex”. In: Progress in Brain Research 114, pp. 359–381.

Pastor, Angel M., Blas Torres, José M. Delgado-Garcia, and Robert Baker (1991). “Discharge characteristics of medial rectus and abducens motoneurons in the goldfish”. In: Journal of Neurophysiology 66.6, pp. 2125–2140.

Pereira, Ulises and Nicolas Brunel (2018). “Attractor Dynamics in Networks with Learning Rules Inferred from In Vivo Data”. In: Neuron 99.1, 227–238.e4.

Pfeifer, Rolf, Max Lungarella, and Fumiya Iida (2007). “Self-organization, embodiment, and biologically inspired robotics”. In: Science 318.5853, pp. 1088–1093.

Sanders, Honi, Michiel Berends, Guy Major, Mark S. Goldman, and John E. Lisman (2013). “NMDA and GABAB (KIR) conductances: the “perfect couple” for bistability.” In: The Journal of Neuroscience 33.2, pp. 424–9.

Seung, H. Sebastian (1996). “How the brain keeps the eyes still.” In: Proceedings of the National Academy of Sciences 93.23, pp. 13339–13344.

Seung, H. Sebastian, Daniel D. Lee, Ben Y. Reis, and David W. Tank (2000). “Stability of the memory of eye position in a recurrent network of conductance-based model neurons”. In: Neuron 26.1, pp. 259–271.

Steinbach, Martin J. (1987). “Proprioceptive knowledge of eye position”. In: Vision Research 27.10, pp. 1737–1744.

Straka, Hans, James C. Beck, Angel M. Pastor, and Robert Baker (2006). “Nucleus prepositus.” In: Journal of Neurophysiology 96(4), pp. 1963–80.

Tsodyks, Misha V. and Henry Markram (1997). “The neural code between neocortical pyramidal neurons depends on neurotransmitter release probability”. In: Proceedings of the National Academy of Sciences 94.2, pp. 719–723.

Urbanczik, Robert and Walter Senn (2014). “Learning by the dendritic prediction of somatic spiking”. In: Neuron 81.3, pp. 521–528.

Vishwanathan, Ashwin, Kayvon Daie, Alexandro D. Ramirez, Jeff W. Lichtman, Emre R.F. Aksay, and H. Sebastian Seung (2017). “Electron Microscopic Reconstruction of Functionally Identified Cells in a Neural Integrator”. In: Current Biology 27.14, 2137–2147.e3.

Wolpert, Daniel M., R. Chris Miall, and Kawato Mitsuo (1998). “Internal models in the cerebellum”. In: Trends in Cognitive Sciences – 2.9, pp. 338–347.

